# Characterization and modulation of human insulin degrading enzyme conformational dynamics to control enzyme activity

**DOI:** 10.1101/2024.12.30.630732

**Authors:** Jordan M. Mancl, Wenguang G. Liang, Nicholas L. Bayhi, Hui Wei, William Budell, Joshua H. Mendez, Tobin R. Sosnick, Bridget Carragher, Clinton S. Potter, Wei-Jen Tang

## Abstract

Insulin degrading enzyme (IDE) is a dimeric M16A zinc metalloprotease that degrades amyloidogenic peptides diverse in shape and sequence, including insulin and amyloid-β, to prevent toxic amyloid fibril formation. IDE has a hollow catalytic chamber formed by two ∼55 kDa N- and C-domains (IDE-N and IDE-C, respectively), in which peptides bind, unfold, and are repositioned for proteolysis. IDE is known to transition between a closed state, poised for catalysis, and an open state, able to release cleavage products and bind a new substrate. Here, we present six cryo-EM structures of the IDE dimer at 3.0-5.1 Å resolution, obtained in the presence of a sub-saturating concentration of insulin. Combining cryo-EM heterogeneity analysis with all-atom molecular dynamics (MD) simulations, we identified the structural basis and key residues for IDE conformational dynamics that were not previously revealed by IDE static structures. Notably R668 serves as a molecular latch mediating the open-close transition and facilitates key protein motions through charge-swapping interactions at the IDE-N/C interface. Our small-angle X-ray scattering analysis and enzymatic assays of an R668A mutant indicate a profound alteration of conformational dynamics and catalytic activity. By integrating coarse-grained MD simulations, our analysis reveals that IDE unfolds its substrates through the coordinated motion between IDE-N and IDE-C, as well as β-sheet formation between IDE and insulin. Additionally, our time-resolved cryo-EM analysis uncovers IDE allostery within the IDE dimer. Collectively, our findings demonstrate the strength of combining experimental and computational approaches to probe protein dynamics and pave the way for developing substrate-specific modulators of IDE activity.

## Introduction

<u>Prot</u>ein home<u>ostasis</u> (a.k.a. proteostasis) is maintained by three primary mechanisms: chaperones, ubiquitination/proteasome, and autophagy (Hipp *et al*, 2019; Lopez-Otin *et al*, 2013). Disruptions in proteostasis can lead to the accumulation of amyloid fibrils and subsequent human diseases (Chiti & Dobson, 2006; Chiti & Dobson, 2017; Greenwald & Riek, 2010). As a result, many proteases have evolved to specifically target amyloidogenic peptides (Malito *et al*, 2008; Saido & Leissring, 2012). Among those, the cryptidase family, which includes the M16 metalloproteases insulin <u>d</u>egrading <u>e</u>nzyme (IDE) use an internal catalytic chamber, or “crypt”, to capture and selectively degrade the monomeric form of amyloid peptides to control the formation of amyloid fibrils (Liang *et al*, 2022; Malito *et al*., 2008). IDE effectively degrades various bioactive peptides, including amyloid-β (Aβ), a peptide associated with the progress of Alzheimer’s disease, and three blood glucose-regulating hormones: insulin, amylin, and glucagon (Tang, 2016). Consequently, defects in IDE alter the progression of type 2 diabetes mellitus and Alzheimer’s disease in animal models and are linked to these diseases in humans (Bertram *et al*, 2000; Bertram *et al*, 2007; Farris *et al*, 2003; Farris *et al*, 2004; Kim *et al*, 2007; Sladek *et al*, 2007; Tang, 2016; Tanzi & Bertram, 2005; Zeggini *et al*, 2007). IDE is a promising therapeutic target, as its inhibition improves glucose tolerance, yet progress has been hampered by the contrary actions of its diverse substrate pool (Maianti *et al*, 2014; Maianti *et al*, 2019; Tang, 2016). For example, IDE overexpression has been shown to reduce Aβ loads, in mice, however, this reduction is accompanied by hyperglycemia from a similar reduction in insulin levels (Leissring *et al*, 2003).

IDE-mediated substrate recognition and degradation is a complex process. IDE is capable of degrading substrates with diverse sequences and structures, such as Aβ and the three glucose-regulating hormones: insulin, amylin, and glucagon (Malito *et al*., 2008; Tang, 2016). Despite this diversity, IDE exhibits high selectivity—for example, it preferentially degrades insulin and atrial natriuretic peptide over insulin-like growth factor-1 and brain natriuretic peptide, respectively (Malito *et al*., 2008). The presence of ATP, inositol phosphate, or specific mutations can markedly enhance the catalytic rate for small peptide substrates like bradykinin, but does not significantly affect the degradation of larger substrates such as insulin and Aβ (Im *et al*, 2007; McCord *et al*, 2013; Song *et al*, 2017; Song *et al*, 2004). Additionally, the binding location of IDE inhibitors within the catalytic chamber can influence substrate preference, making IDE more likely to degrade one substrate over another (Charton *et al*, 2014; Maianti *et al*., 2019). This complexity arises partly from the interplay between the unfoldase and protease activities of IDE, which are essential for degrading diverse substrates (Malito *et al*., 2008; Tang, 2016; Zhang *et al*, 2018). IDE relies on its unique large catalytic chamber and associated conformational dynamics to unfold and degrade amyloidogenic substrates. A deeper understanding of the molecular mechanisms by which IDE utilizes its catalytic chamber for substrate unfolding and degradation is crucial for effectively harnessing this enzyme in future therapeutic development.

IDE is a dimeric 110 kDa metalloprotease. Each protomer is comprised of four homologous subdomains organized into two ∼55 kDa domains, IDE-N and IDE-C, connected by a short flexible linker (Li *et al*, 2006; Shen *et al*, 2006) (Fig. 1A). Our previous cryo-EM analysis revealed the major conformational states of IDE. In the absence of substrate, each protomer was found to adopt an open (O) or partial open (pO) state which are primarily differentiated by the displacement of IDE-N relative to IDE-C, the combination of which led to three dimeric conformations (O/O, O/pO, pO/pO) (Zhang *et al*., 2018). It was observed that substrate binding induced both protomers of IDE to fully close, referred to as the partial closed (pC, pC/pC for the dimer) state, as the cryo-EM closed state was found to be slightly more open than previously solved crystal structures, likely influenced by the constraints of the crystal lattice (McCord *et al*., 2013; Shen *et al*., 2006). Only the O state permits substrate access to the catalytic chamber, yet the catalytic cleft is stabilized only in the pC state with substrate bound, requiring IDE to undergo a substantial open-close transition during catalysis, the specifics of which are unknown (Zhang *et al*., 2018).

**Figure 1:**
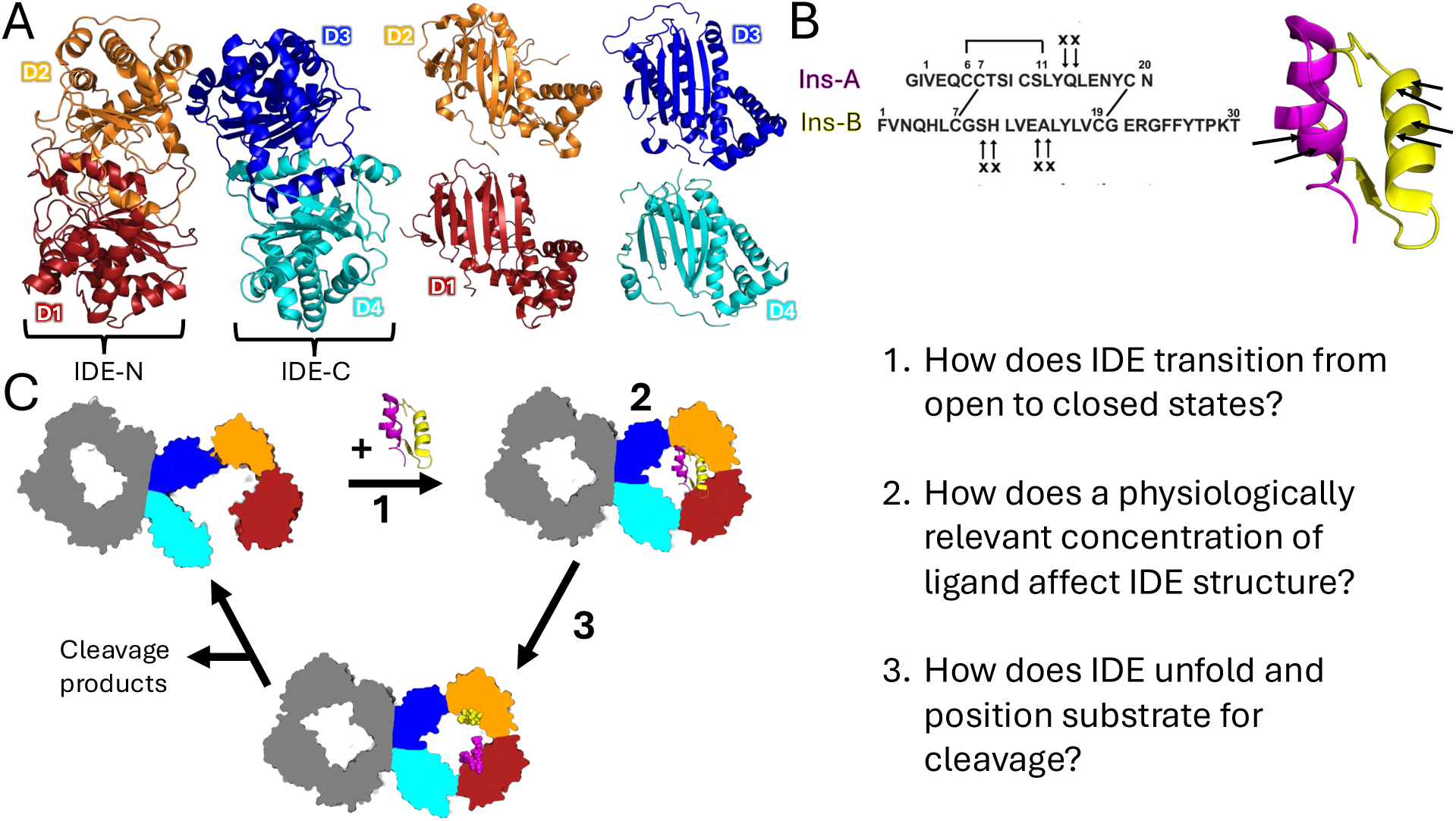
Overview of structure and function of IDE. (A) Overall structure of IDE. IDE is comprised of four structurally homologous domains: D1 (red, residues 43-285), D2 (orange, residues 286-515), D3 (blue, residues 545-768), and D4 (cyan, residues 769-1,016). The domains are arranged in two roughly hemispherical regions that enclose the catalytic chamber referred to as IDE-N (comprised of the D1 and D2 domains) and IDE-C (comprising the D3 and D4 domains), which are joined by a linker region (residues 516-544). (B) Primary sequence and overall structure of insulin. Insulin cleavage by IDE as revealed by mass spectrometry (Manolopoulou *et al*., 2009). Cleavage sites are marked in arrows, which occur within α-helical regions and requires the peptide to be unfolded prior to cleavage. (C) Overview of the IDE catalytic cycle with key questions addressed in this study.

Biochemical and structural analyses indicate that many IDE substrates, such as insulin, must first be unfolded prior to cleavage (Fig. 1B, C). The conformational dynamics of IDE are thought to play a central role in substrate recognition and degradation, yet several mechanistic details underlying the unfoldase activity of IDE remain unresolved (Fig. 1). We hypothesize that the unfoldase and protease activities of IDE are regulated by the relative motions of the IDE-N and IDE-C domains, their interactions with substrates, and allosteric communication between subunits within the IDE dimer. To investigate this, we solved cryo-EM structures of IDE bound to insulin at a 2:1 IDE:insulin ratio, as well as structures captured following rapid mixing of IDE with insulin under functionally relevant conditions. Utilizing newly developed computational methods, we analyzed particle heterogeneity in these structures to infer the conformational dynamics of IDE. By integrating these experimental data with all-atom and coarse-grained molecular dynamics (MD) simulations, we identified and modulated specific molecular interactions to manipulate key conformational dynamics of IDE, ultimately enhancing our understanding how to control its enzymatic activity in vitro.

## Results

### Cryo-EM reveals new IDE structures in the presence of a sub-saturating concentration of insulin

We have previously used cryo-EM to solve the structure of IDE in the presence of a 5-fold molar excess of insulin, resulting in a dominant pC/pC state with both substrate binding chambers yielding density corresponding to unfolded insulin (Zhang *et al*., 2018). To explore the mechanism of IDE-directed substrate unfolding under more physiologically relevant conditions, we investigated IDE-insulin interactions with an IDE:insulin molar ratio of 2:1. We collected a dataset of ∼7,600 micrographs on a Titan Krios operated at 300 keV, from which 7.2 million particles were picked for processing in RELION. Following an established workflow, and processing the data with C1 symmetry, we generated five structures: three previously observed structures with improved resolution (pC/pC, O/O, and O/pO at 3.0 Å, 3.8 Å, and 4.1 Å resolution) and two novel states (O/pC and pO/pC at 3.4 Å and 3.3 Å resolution, respectively) (Fig. 2A, Figure 2 – figure supplement 1, Supplementary file 1). Interestingly, the pC/pC was still found to be dominant, despite IDE and insulin being present at a 2:1 molar ratio, consistent with the allostery of IDE (McCord *et al*., 2013; Zhang *et al*., 2018).

**Figure 2:**
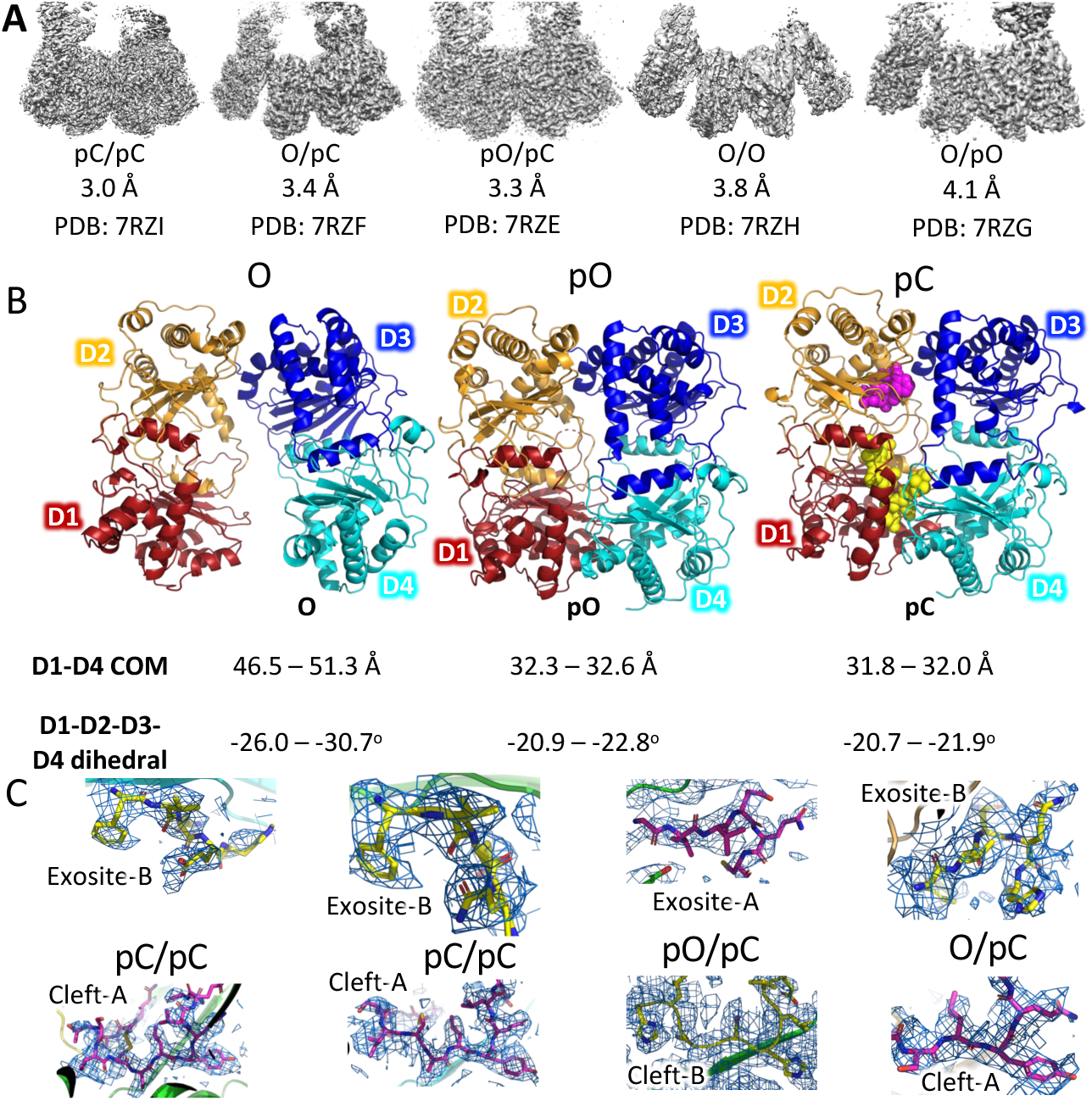
Cryo-EM structures. **(A)** Overview of the cryo-EM structures. See figure S1 for processing details. **(B)** Comparison of the open (O), partial open (pO), and partial closed (pC) subunit states present in our cryo-EM structures with domain organization. The distance between the D1 and D4 domain centers-of-mass (D1-D4 COM) along with the dihedral angle formed by the D1-D2-D3-D4 domain centers-of-mass (D1-D2-D3-D4 dihedral) described in Zhang et al. (21) and depicted in Figure 3 – figure supplement 1 were used as biologically important criteria to quantify observed conformations. **(C)** Insulin density and corresponding model in our cryo-EM structures. Both the A chain (magenta) and B chain (yellow) can fit the density in the exosite and catalytic cleft.

The individual protomers of IDE adopt the same three conformations: O, pO, and pC states, reported previously but there are significant differences in the O and pO states (Fig. 2B) (Zhang *et al*., 2018). While the pC/pC state is nearly identical to the two states reported previously (RMSD of ∼0.6 Å; PDB ID: 6BFC, 6B3Q) (Zhang *et al*., 2018), the O and pO states appear more closed in the presence of insulin, suggesting that the presence of insulin promotes the open to closed transition. Specifically, the pO and O subunits are 3-5 Å more closed based on the distance between the centers-of-mass (COM) of the IDE D1 and D4 subdomains, while the dihedral angle formed by the COM of the D1-D2-D3-D4 subdomains is reduced by 5-10° (Fig. 2B). Alignment of the new O/O and O/pO states to their previously solved counterparts reveals little difference, with a global RMSD of 1.316 Å (O/O, PDB ID: 6B7Y) and 1.405 Å (O/pO, PDB ID: 6BF8), respectively (Zhang *et al*., 2018). The primary source of variation between the different states remains the degree of opening between the two domains of each subunit, as a global alignment of IDE-N and IDE-C across all structures reveal the domains primarily act as rigid bodies (RMSD <1.4 Å). Consistent with previous observations, the door region, which contains the catalytic zinc binding site, exhibited higher B-factors in the O states than in either the pO or pC state (Zhang *et al*., 2018) (Figure 2 – figure supplement 2).

The cryo-EM structures presented here contain four subunits that have clear density present at the catalytic cleft and exosite indicative of bound substrate, thus adopting the pC state. Previous structures have modeled the same chain of insulin in both the exosite and catalytic cleft, but the best models we could build into our density suggest that the chain bound to the exosite is not the same chain positioned for cleavage at the catalytic cleft (Fig. 2C). The preponderance of data suggests it to be unlikely that insulin adopts a preferred orientation within the catalytic chamber, rather the catalytic chamber can accommodate insulin in four different orientations, with either the A or B chain bound to the exosite and catalytic cleft in *cis* and *trans* orientations. This promiscuity of binding has been theorized as a reason for why there is no chain preference for the initial insulin cleavage event (Manolopoulou *et al*, 2009). Crystal structures of insulin are highly compact and globular, with little spatial separation between the N-and C-termini of each chain, and the IDE cleavage sites are located within α-helices (Baker *et al*, 1988). In its crystallized state, insulin cannot interact with both the exosite and catalytic cleft unless it unfolds, at least partially. Recent work used MD simulations to study the conformational dynamics of insulin in solution and identified several major “elements of disorder” to describe the partially unfolded intermediate structures they observed (Busto-Moner *et al*, 2021). We docked these structures of insulin into the closed catalytic chamber of IDE and found that the chamber was able to easily accommodate the positioning of either chain near the exosite and catalytic cleft in *cis* and *trans* orientations (Figure 2 – figure supplement 3).

### Heterogeneity analysis reveals 2 dominant components of structural variability with the IDE cryo-EM particle population

Numerous structures of IDE exist with subunits adopting either an open or closed conformation, yet we lack an understanding of how IDE transitions from an open to a closed state. The simplest explanation of this transition, based on structural data, would be a direct, rigid body translation of IDE-N relative to IDE-C, and this is the predominant model within the field (Shen *et al*., 2006; Zhang *et al*., 2018). To better understand what this transition would look like, we measured and plotted the changes in the D1-D4 COM distance versus the changes in the D1-D2-D3-D4 COM dihedral angle for all available cryo-EM structures of IDE (Figure 3 – figure supplement 1). We found that in the absence of substrate, the transition pathway produced a linear relationship between states. However, when our structures generated in the presence of insulin were analyzed, the linear relationship no longer held (Figure 3 – figure supplement 1), suggesting that the simple linear transition typically presumed for the open-close transition between distinct IDE states does not accurately depict the complexity of IDE dynamics.

Recently, several approaches have been developed to understand the conformational heterogeneity present within cryo-EM data. We employed multibody analysis in RELION and 3D variability analysis (3DVA) in cryoSPARC to investigate the conformational heterogeneity within our particle populations (Nakane *et al*, 2018; Punjani & Fleet, 2021). Of these, the range of motion predicted by multibody analysis most closely matched the reported structures while 3DVA exhibited consistent results across a smaller magnitude of transition (Figure 3 – figure supplements 1-3). Therefore, we focused on multibody analysis below. Multibody analysis models the structural heterogeneity in our data as the result of motions of independent, user-defined rigid bodies. As discussed above, the primary source of structural variation in our structures is the positioning of IDE-N relative to IDE-C. We see little structural changes within the domains, suggesting that IDE domains behave as three rigid bodies, IDE-N in two different subunits within IDE dimer (IDE-N A, IDE-N B) and the dimeric IDE-C (IDE C/C). This is consistent with the structural analysis of M16 family of metalloproteases (Liang *et al*., 2022). This assertion is further supported by our all-atom molecular dynamics (MD) simulations. While we observed small yet noticeable conformational changes within IDE-N A, IDE-N B, and IDE-C/C, the overall RMSD of entire IDE dimer is significantly higher than those three individual units (Fig. 3A). Multibody analysis defines the principal components of variance along discrete eigenvectors, which we interpret as proxies for the dominant components of molecular motion. We quantified the change in D1-D4 COM distance and D1-D2-D3-D4 COM dihedral of representative structures along the heterogeneity gradients of the top 9 eigenvectors for each of our IDE structures, representing ∼70-90% of the total structural heterogeneity per structure. The results indicated that the particles comprising our cryo-EM structures sampled a significantly greater conformational space than would be expected from analysis of the ensemble structures alone (Figure 3 - figure supplement 2). Most intriguingly, the conformational changes displayed an unexpectedly high degree of change in the D1-D2-D3-D4 COM dihedral angle, indicating a significant rotation of IDE-N relative to IDE-C (Fig. 3B,C, Figure 3 – figure supplement 2).

**Figure 3:**
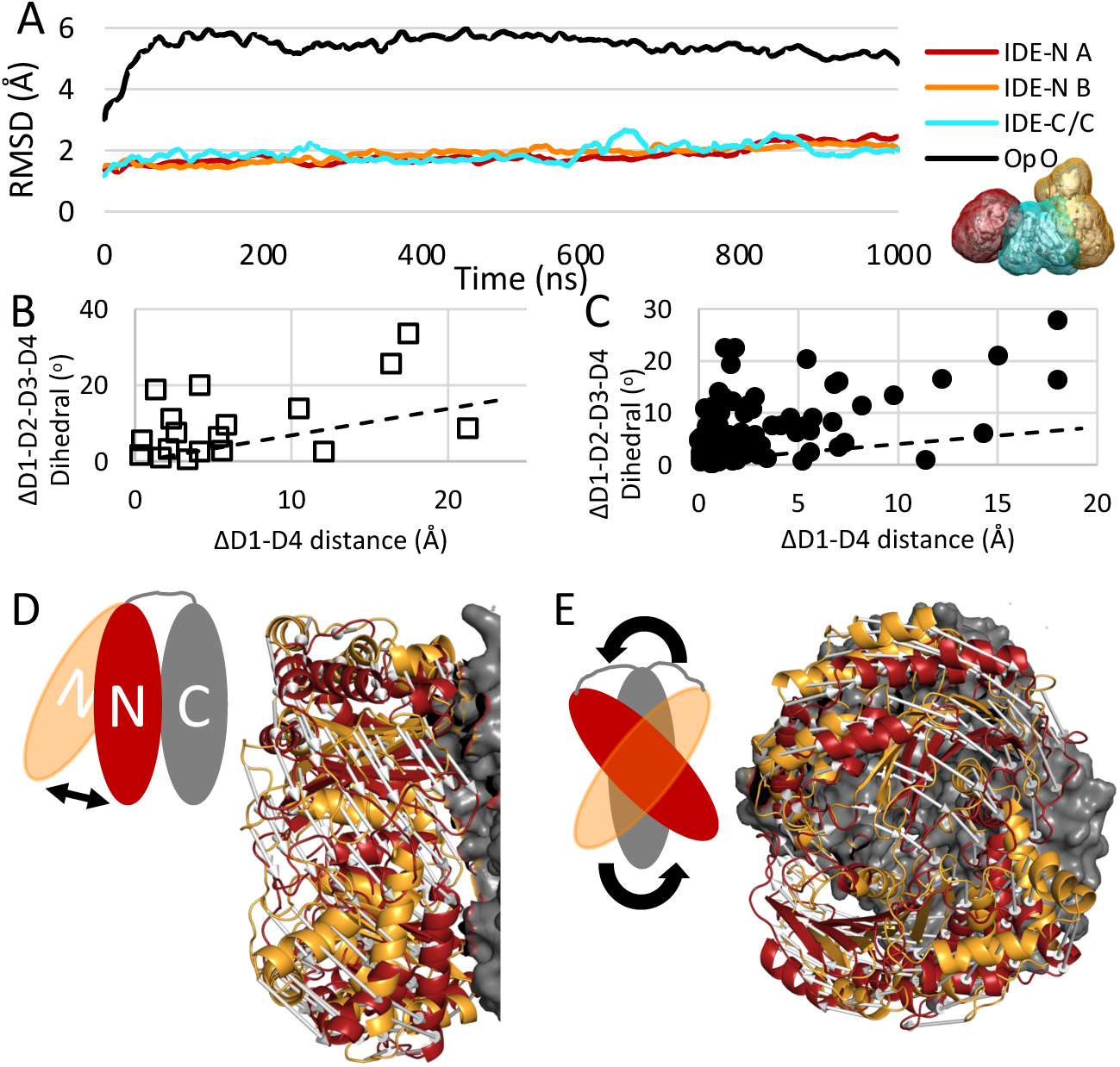
Conformational dynamics of IDE implied by structural heterogeneity. **(A)** All-atom MD simulations analysis. The primary source of structural variance (RMSD) results from the IDE-N moving against IDE-C as a rigid body. Rigid bodies were defined as colored for multibody refinement in RELION. **(B-C)** Multibody analysis. The range of conformational variance described by the top principal component vectors displays an unexpectedly high degree of rotational motion, as measured by the change in D1-D2-D3-D4 dihedral angle across the gradient of structural heterogeneity for each vector, in both the absence **(B)** and presence **(C)** of insulin compared to the expected open-close transition pathway predicted from a linear interpolation of the experimentally determined structures of IDE (dashed line, Figure 3 – figure supplement 1). Two dominant components of structural variance are revealed from multibody analysis: **(D)** where IDE-N swings relative to IDE-N about the inter-domain linker, and **(E)** where IDE-N rotates against IDE-C. Starting (orange) and ending (red) states of IDE-N shown with pathway depicted by arrows. IDE-C shown as gray surface.

Integration of the multibody results across all states of IDE suggest two dominant components of structural variance. First is a translation-dominant conformational change wherein IDE-N swings toward or away from IDE-C as if mediated by a hinge formed by the interdomain linker region of IDE (Fig. 3D, Figure 3 - video 1). This conformational change closely resembles our understanding of the IDE open-closed transition inferred from analysis of the ensemble structures. The second component is a rotation-dominant conformational change wherein IDE-N rotates orthogonal to the plane of the dimer as if it were being screwed into or ground against IDE-C (Fig. 3E, Figure 3 – video 2). Importantly, these dominant components of structural variance were also observed when the particle populations were analyzed with 3DVA in cryoSPARC, albeit over a smaller magnitude of conformational change (Figure 3 – figure supplement 3). These dominant components also correlate well with the lowest frequency modes revealed by normal mode analysis (Figure 3 – videos 3, 4). Interestingly, while the presence of insulin was found to substantially influence the consensus reconstructions of IDE, we found that the presence or absence of insulin had no significant effect on the principal components of structural variance for IDE, consistent with the current understanding that enzyme conformational changes are “hardwired” into the structure (Bahar *et al*, 2007; Bahar & Rader, 2005).

### R668 plays a key role in the IDE open/close transition in all-atom MD simulations

Analysis of the particle heterogeneity comprising cryo-EM structures provides excellent information about the conformational space sampled by the protein of interest. However, such analysis lacks a temporal component, and thus only implies motions. To overcome this barrier and examine how the conformation of IDE changes over time, we performed six replicate all-atom MD simulations with explicit solvent. Simulations were initialized from the O/pO state (PDB: 7RZG), with the missing loops modeled in to generate continuous peptide chains and run for one microsecond each. By starting with the O/pO structure, we investigated the dynamics of both the open and closed subunits simultaneously. We observed that the conformational space sampled by our simulations correlated well with the conformational space sampled by our cryo-EM particle population, as revealed by multibody analysis (Figure 4 – figure supplement 1). In five of our six simulations, the open subunit closed quickly, typically in <200 ns (Fig. 4A). Analysis of the open-close transition revealed two key findings. First, the open subunits did not close to a singular consensus structure. We define closing as a subunit reaching a similar D1-D4 COM distance as the pO or pC states, but we observed distinct differences among the subunits indicative of IDE-N rotation relative to IDE-C. These differences are in line with our second key finding, that the conformational changes associated with the open-close transition did not follow the same general pathway among the simulations, as we observed a high degree of variability upon closing (Fig. 4B).

**Figure 4:**
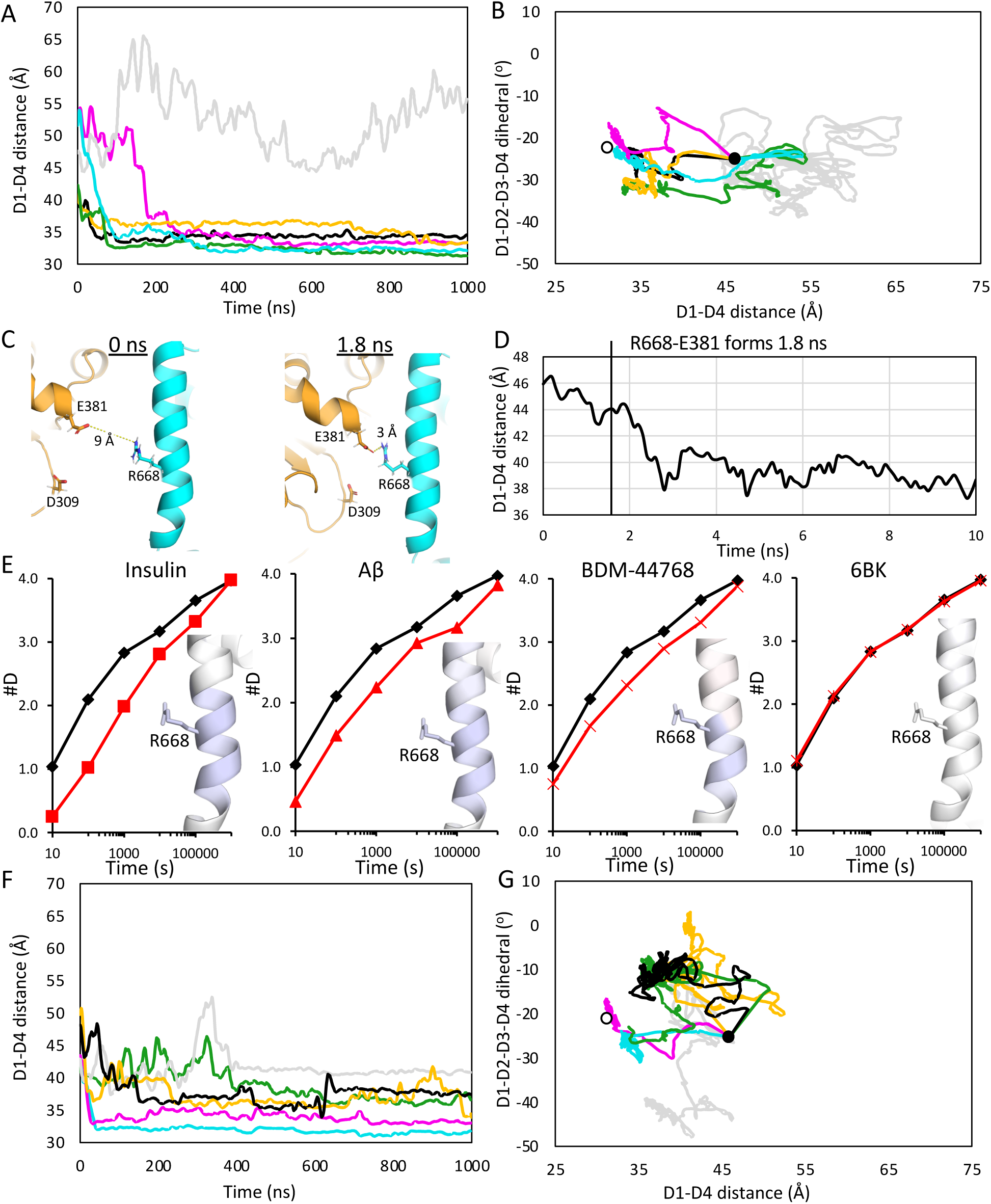
All-atom MD reveals a molecular basis for IDE conformational dynamics. **(A)** Measurements of the O subunit D1-D4 distance over the course of six separate microsecond long all-atom MD simulations of WT IDE. Of which, the open subunit closed in 5 of the 6 simulations. **(B)** Plot of the O subunit D1-D4 distance vs the D1-D2-D3-D4 COM dihedral angle over the course of the simulation of WT IDE. The open subunits displayed a variety of closing pathways and did not close to a consensus structure. Starting structure shown as black dot, pO structure shown as white dot. **(C)** R668 acts as a guidepost residue, rapidly interacting with D309 or E381. Formation of this interaction is associated with rapid closing, as measured by a decrease in D1-D4 distance **(D). (E)** Hydrogen-deuterium exchange mass spectrometry highlights the importance of R668 in mediating the open-close transition. In the presence of insulin (panel 1, red), Aβ (panel 2, red), and BDM-44768 (panel 3, red), all of which promote IDE closing, the peptide containing R668 shows reduced deuterium exchange relative to apo-IDE (black), yet in the presence of 6bk (panel 4, red), which does not promote closing, there is no difference in the exchange rates for the R668 containing peptide relative to apo-IDE (black). Helix containing R668 colored by red (increase) – white (no change) – blue (decrease) gradient depicting the degree of deuterium exchange relative to apo-IDE. **(F)** Measurements of the O subunit D1-D4 distance over the course of six separate microsecond long all-atom MD simulations of IDE R668A. **(G)** Plot of the O subunit D1-D4 distance vs the D1-D2-D3-D4 COM dihedral angle over the course of the simulation of IDE R668A. The six separate microsecond long simulations indicate that an R668A mutation significantly alters the closing dynamics of IDE **(F)** and increases the rotational motion **(G)** relative to WT (panels **A** and **B** respectively). Starting structure shown as black dot, pO structure shown as white dot.

Noticeably, R668 consistently stood out as one of the first IDE-C residues to form new interactions with IDE-N, either D309 or E381 despite the path variation among our MD simulations (Fig. 4C). In most simulations, formation of this interaction preceded a large decrease in the D1-D4 COM distance (Fig. 4D, Figure 4 – figure supplement 2). Interestingly, these R668 interactions are not present in the ensemble cryo-EM or crystal structures of IDE. We hypothesize that the biochemical properties and spatial localization of R668 enable it to essentially reach out and grab onto the N-domain, initiating the formation of a complex hydrogen bonding network that drives the open-close transition towards completion. Previously published hydrogen-deuterium exchange mass spectrometry (HDX-MS) data indicates that R668 is stabilized under conditions that promote IDE closing, suggesting that R668 may function as a molecular latch (Fig. 4E) (Zhang *et al*., 2018).

To further probe the importance of R668, we ran 6 additional simulations with IDE carrying a R668 to alanine (R668A) point mutation under identical conditions as described previously. These simulations displayed significantly altered dynamics when compared to the wild-type simulations. Notably, the R668A mutant was found to close more slowly, and many simulations did not reach the D1-D4 COM distance typified by a closed state structure, instead stabilizing with a significantly larger D1-D4 COM distance than we had previously observed in our wild type (WT) simulations or our experimental structures (Fig. 4F). We also observed that the R668A mutant preferentially sampled a subset of conformational space rarely explored in our WT simulations and displayed significantly greater variation in the D1-D2-D3-D4 dihedral angle (Fig. 4G). As a result of this increased rotational exploration, several of our R668A simulations closed to a conformation where IDE-N is offset and rotated relative to IDE-C when compared to the experimentally determined structures (Figure 4 – figure supplement 3). This altered conformation results in a structure where IDE-N and IDE-C are fully engaged, but the offset produces a deceivingly high D1-D4 COM distance. With this in mind, we sought to better understand the conformational dynamics of IDE in the closed state.

### R668A modulates IDE conformational dynamics *in vitro*

The R668A mutation was found to significantly alter the conformational dynamics of the IDE open-close transition in our simulations (Fig. 4F, G), so we expressed and purified the R668A mutant to determine if the changes predicted from our MD simulations resulted in altered enzymatic activity relative to WT. Previous work has established that IDE adopts a dominant O/pO state in solution (Zhang *et al*., 2018). Consistent with our MD simulations, size-exclusion chromatography revealed that the R668A construct eluted slightly earlier than WT IDE, suggesting that the R668A mutation induces a greater proportion of molecules to adopt a more open conformation, possibly shifting to a dominant O/O state (Fig. 5A). Next, we compared the enzymatic activity of the R668A and WT constructs using the fluorogenic substrate (7-methoxycourmarin-4-yl)acetyl-RPPGFSAFK(2,4-dinitrophenyl)-OH (substrate V), a bradykinin mimetic which has previously been used to characterize the enzymatic activity of IDE and related enzymes (King *et al*, 2014; McCord *et al*., 2013). We found that WT IDE has activity consistent with previously published data (McCord *et al*., 2013) and the R668A mutation produced a ∼5-fold decrease in activity (Fig. 5B). ATP is known to promote IDE degradation of small substrates, i.e. bradykinin, but not larger substrates, i.e. insulin (33). Consistent with this observation, the addition of ATP increased the activity of both WT and R668A constructs (Fig. 5B). We also tested if the R668A mutation would alter the degradation of larger, well-folded substrates, using insulin in a direct competition assay. We found that, after accounting for the previously observed 5x decrease in activity, insulin yielded an apparent K_i_ of ∼8 nM for WT IDE, but this value was reduced to ∼52 nM for the R668A construct (Fig. 5C). When Aβ(1-40), an unstructured substrate, was tested in the same competition assay, we observed an apparent K_i_ of ∼688 nM for WT IDE, yet the R668A mutant did not exhibit a dose-dependent response to the presence of Aβ and the data could not be fit to any kinetic equation (Fig. 5D). We then performed size exclusion chromatography-coupled small-angle X-ray scattering (SEC-SAXS) for both constructs to ascertain if the observed differences in enzymatic activity could be explained by altered biophysical properties. The WT scattering profile produced an R_g_ of 49.4 +/-0.9 Å, consistent with previous SEC-SAXS results (Fig. 5E, F) (Zhang *et al*., 2018). The R668A scattering profile produced an R_g_ of 54.7 +/- 0.4 Å, consistent with our SEC data that R668A mutant has slightly larger hydrodynamic radius (Fig. 5A). Together, our data indicates that R668A mutation can profoundly affect the conformational dynamics governing the open-closed transition, and that altered conformational dynamics can alter enzymatic activity in a substrate-dependent manner.

**Figure 5:**
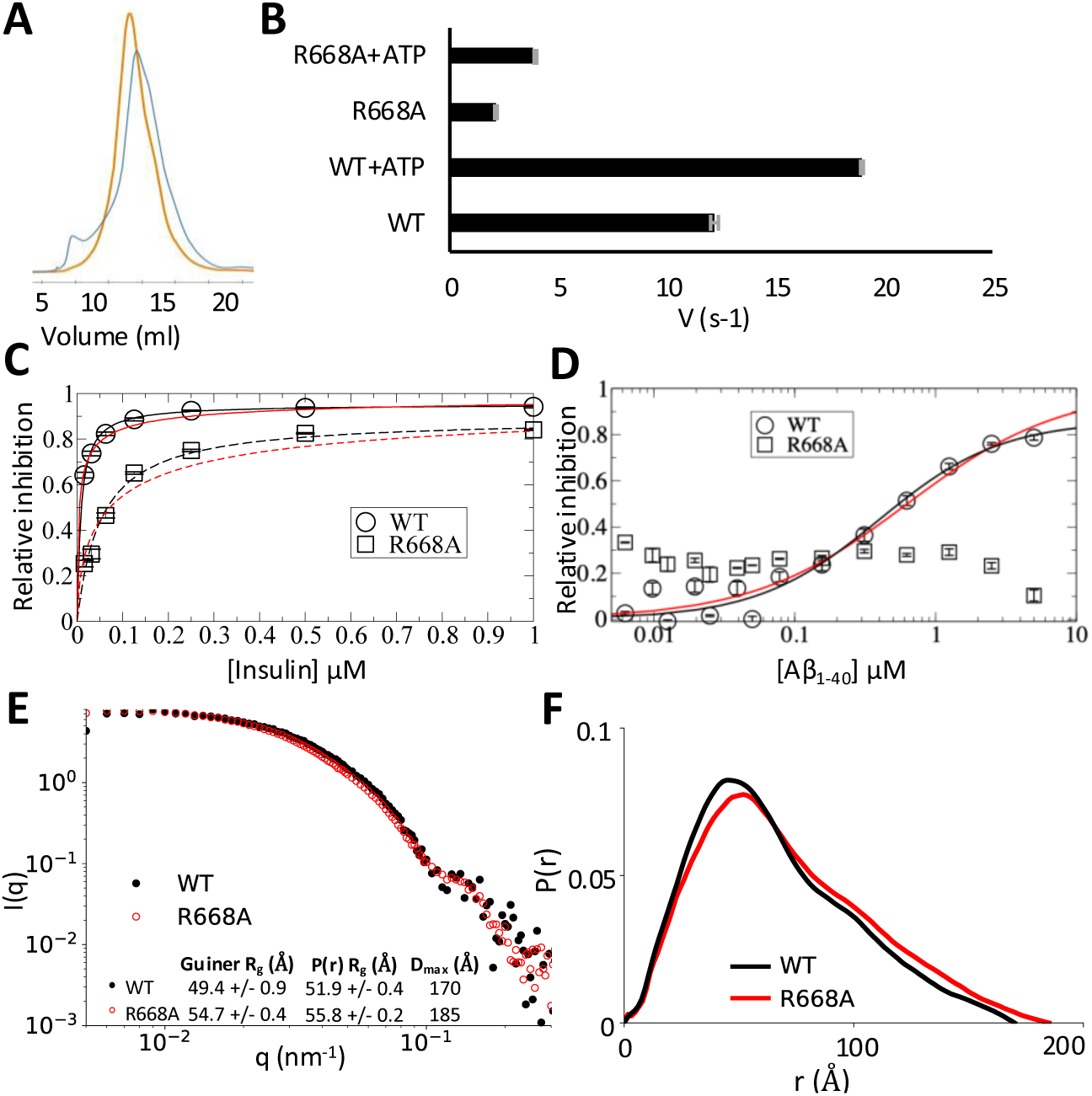
R668A alters IDE activity *in vitro.* **(A)** Elution profile of WT IDE (blue) compared to the R668A mutant (orange) from a S200 SEC column. (**B**) Degradation of the fluorescent substrate MCA-RPPGFSAFK(Dnp) by WT IDE and the R668A construct in the presence and absence of ATP. Data represents the average initial velocities of three replicates performed at a protein concentration of 3.125 nM. Error bars (gray) represent the standard error. **(C)** Inhibition of MCA-RPPGFSAFK(Dnp) degradation by WT IDE (circles, solid fit lines) and IDE R668A (squares, dashed fit lines) in the presence of varying amounts of insulin. Data was fit to the Michaelis-Menten (black) and Hill equations (red). Relevant parameters, Michaelis-Menten: WT: χ^2^=0.001, V_max_=0.951, K_i_=8.3 nM; R668A: χ^2^=0.005, V_max_=0.892, K_i_=52 nM; Hill: WT: χ^2^=0.009, n=0.55, K_i_=51 nM; R668A: χ^2^=0.055, n=0.61, K_i_=198 nM. Error bars represent standard error, data points represent the average of three technical replicates. **(D)** Inhibition of MCA-RPPGFSAFK(Dnp) degradation by WT IDE (circles, solid fit lines) and IDE R668A (squares, dashed fit lines) in the presence of varying amounts of Aβ_1-40_. WT data was fit to the Michaelis-Menten (black) and Hill equations (red), R668A data could not be fit to either equation, indicating that the mutation confers substrate-specific altered enzyme kinetics. Relevant WT parameters, Michaelis-Menten: χ_2_=0.066, V_max_=0.823, K_i_=353 nM; Hill: χ^2^=0.08, n=0.72, K_i_=688 nM. Error bars represent standard error, data points represent the average of three technical replicates **(E)** SEC-SAXS profile of WT (black) and R668A (red) constructs with Rg values calculated by both the Guinier and Porod methods along with Dmax derived from the P(r) function **(F)**.

### Charge-swapping at the IDE-N/C interface mediates conformational dynamics in the closed state

The WT pO subunit remained closed throughout all simulations yet was found to initially relax to a slightly more open state within the first 200 ns, roughly the same time frame it took the WT O subunit to close (Fig. 6A) and, despite not opening, demonstrated a range of D1-D2-D3-D4 dihedral variation similar to that of the O subunits (Fig. 6B). Interestingly, the conformational variation within our simulations increased greatly after this relaxation point was reached (Fig. 6A). This relaxed conformational state consistently displayed an altered IDE-N/C interface compared to the experimentally determined structures that was replicated by the open WT subunits upon closing, leading to a nearly symmetric pO/pO state (Figure 6 – figure supplement 1). Biochemically, this relaxed IDE-N/C interface makes more sense than the interface observed experimentally. It has previously been demonstrated that the experimental constraints of crystallographic structure determination force IDE to adopt a closed conformation and determination of the open state can only be accomplished once those restraints are removed, i.e. cryo-EM (Zhang *et al*., 2018). These results suggest that both ensemble methods of structure determination may impact the conformation of IDE, however slightly. The IDE-N/C interface is littered with charged residues, yet few interactions are observed in the ensemble structures (Fig. 6C), whereas our MD simulations reveal a complex hydrogen bonding network (Fig. 6D). We observed an extensive network at the D2-D3 interface, with sporadic patches of interactions between D1 and D4 (Figure 6 – figure supplement 2). Within the D2-D3 network R668 again stands out as a notably key residue. In addition to forming hydrogen bonds with D309 and E381, we observed R668 form *τ-τ* interactions with R311 (Fig. 6E). Arginine-mediated *τ*-interactions have been well characterized in other systems for their ability to stabilize interaction interfaces and facilitate conformational changes (Armstrong *et al*, 2016; Vernon *et al*, 2018).

**Figure 6.**
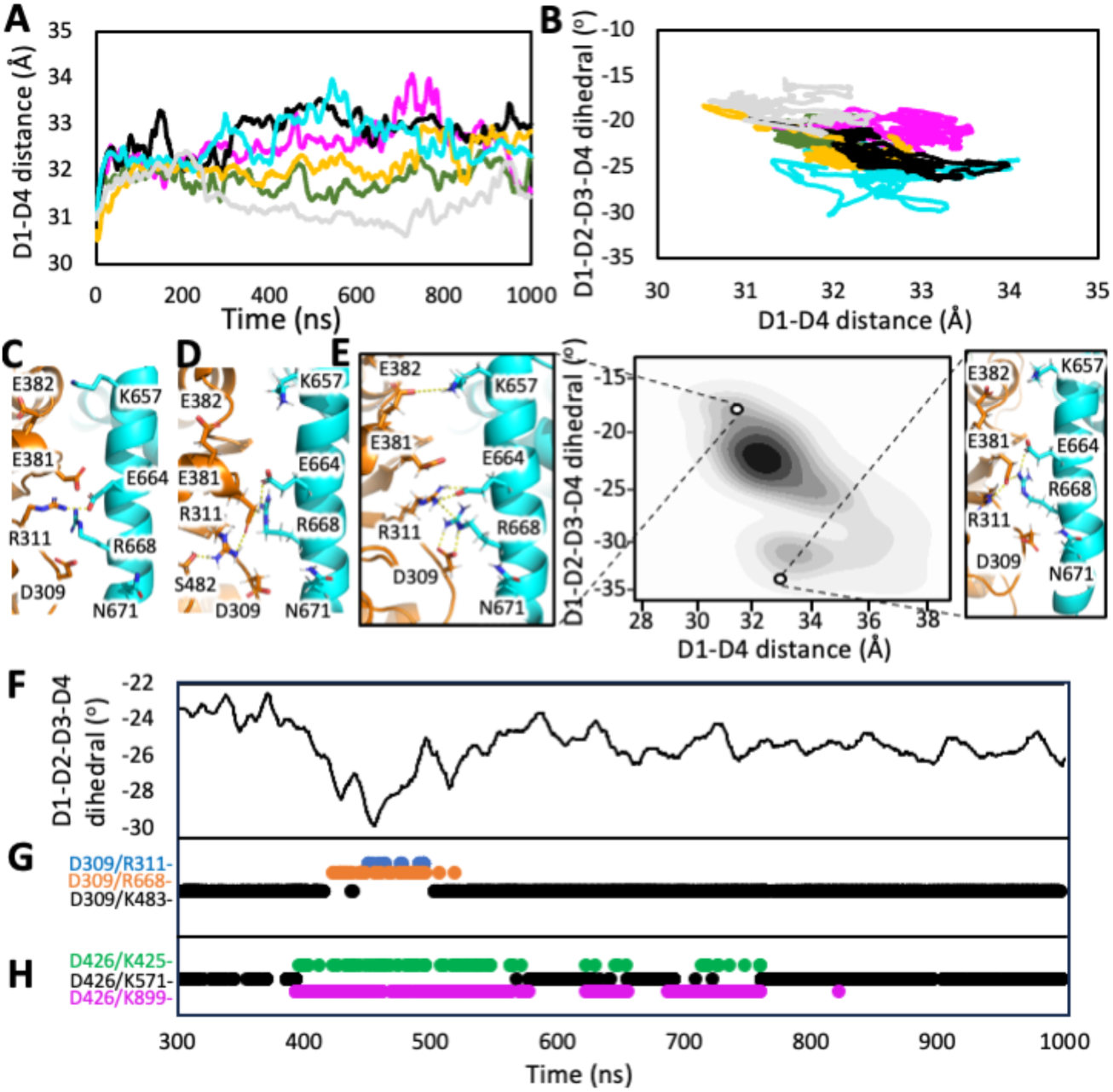
Structural basis of closed state conformational dynamics. **(A)** Measurements of the pO subunit D1-D4 distance over the course of six separate microsecond long all-atom MD simulations of WT IDE. **(B)** Plot of the pO D1-D4 distance vs the D1-D2-D3-D4 COM dihedral angle over the course of the simulation of WT IDE. **(C)** IDE-N/C interface previously solved crystal structures (PDB:2G47 shown) shows side chains are ill-positioned for interaction. **(D)** IDE-N/C interface formed upon open subunit closing in our MD simulations reveals a complex hydrogen bonding network. **(E)** Heat map showing conformational geometries that were preferentially sampled in our MD simulations by the open subunits upon closing. Insets highlight how the IDE-N/C interface changes to permit interdomain motion. **(F)** Plot of the O subunit D1-D2-D3-D4 dihedral angle during a subset of a single WT IDE MD simulation after the open-close transition has been completed. Charge-swapping between residues at the IDE-N/C interface is associated with changes in the D1-D2-D3-D4 dihedral. **(G)** For most of the simulation, D309 interacts with K483 (black), however, this interaction is broken for ∼100 ns, during which D309 instead interacts with R311 (blue) and R668 (orange). **(H)** For most of the simulation, D426 interacts with K571 (black), yet this interaction is periodically broken, and D426 instead interacts with K425 (green) and K899 (magenta). When these events of charge-swapping coincide with D309 charge-swapping **(G)**, they are associated with a large change in the D1-D2-D3-D4 dihedral angle **(F)**. When they occur alone, the effect on D1-D2-D3-D4 dihedral is smaller.

The IDE-N/C interactions we observe in our simulations are not restricted to discrete cognate pairs; rather, we observed that residues within the network periodically swap among several interaction partners. This “charge-swapping” phenomenon allows IDE-N to rotate or slide against IDE-C to adopt multiple conformations and maintain favorable interacting contacts (Fig. 6F). While multiple, concurrent, instances of charge-swapping were associated with rapid changes in the D1-D2-D3-D4 dihedral angle, singular events of charge-swapping did not appear to be significant enough to stimulate large-scale conformational change (Fig. 6G, H). Instead, substantial conformational changes appear to be caused by multiple charge-swapping events occurring in conjunction. This may explain why we observed no noticeable change in the conformational dynamics of the pO state subunits when comparing the WT and R668A constructs. Consurf analysis reveals that most of the residues forming these interaction networks are highly conserved among IDE homologs (Figure 6 – figure supplement 3) (Ben Chorin *et al*, 2020). This indicates that the revealed rotational motion is likely evolutionarily conserved and offers a potential mechanism by which IDE unfolds and repositions bound peptide substrates to degrade amyloid peptides.

### Time-resolved cryo-EM suggests insulin-induced allostery in IDE

To begin understanding of how the interaction between IDE and insulin leads to insulin unfolding and degradation, we first performed all-atom MD simulation of insulin-bound IDE in the closed state. Unfortunately, we did not observe a noticeable structural change of insulin within our microsecond simulations. Previously, we used time-resolved SAXS analysis to determine the time constant for the insulin induced open to closed transition, tau (1, a.k.a., monoexponentially decay) to be ∼100 milliseconds (Zhang *et al*., 2018). However, it has previously been determined that IDE cleaves about two insulin molecules every second (Manolopoulou *et al*., 2009). We employed spotiton to rapidly mix and freeze grids where IDE has been exposed to insulin for 123 milliseconds prior to freezing (Dandey *et al*, 2020). We hoped that such an experimental setup would permit the identification of novel insulin-bound states of IDE and offer insight into the initial binding and unfolding of insulin. Previous work has established that, in the absence of substrate, IDE adopts a dominant O/pO state in solution and the presence of a molar excess of substrate shifts the particle population to a dominant pC/pC state (Zhang *et al*., 2018). Surprisingly, after processing the dataset in Relion, we observed a single conformational state within our rapidly-mixed dataset: the O/O state (Fig. 7A, Figure 7 - figure supplement 1, supplementary file 2). We were unable to resolve the structure beyond moderate resolution (∼5.1 Å) and did not observe unambiguous density indicative of insulin binding within the catalytic chamber, which is unsurprising given our previous observations of promiscuous binding (Fig. 2C). Despite the relatively high particle number in our final reconstruction, we were unable to sub-classify distinctly different conformations of IDE within the particle population as initial classes converged back to the dominant O/O state upon subsequent refinement. This leads us to believe that the obtained O/O state represents the average state of a particle population that exhibits a high degree of continuous structural heterogeneity. The O/O state observed in our rapidly-mixed dataset is distinct from the previously observed O/O states with D1-D4 center of mass distances of 55.1 Å and 46.8 Å matching the largest and smallest, respectively, distances previously observed for individual O state subunits (Fig. 7B). The discrepancy between the “openness” of the two subunits nearly doubles that observed in previous structures, making this O/O state the most asymmetric O/O state yet observed (Fig. 7C). The observation of a dominant O/O state in the presence of insulin under short timescales suggests that the binding of insulin to the O subunit of the apo-dominant O/pO state can induce the pO state to open through an unknown allosteric mechanism to reach the observed dominant O/O state (Fig. 7D). Positive allostery in IDE has previously been identified in biochemical assays, yet we lack a detailed understanding of the underlying mechanism (McCord *et al*., 2013; Song *et al*, 2011).

**Figure 7:**
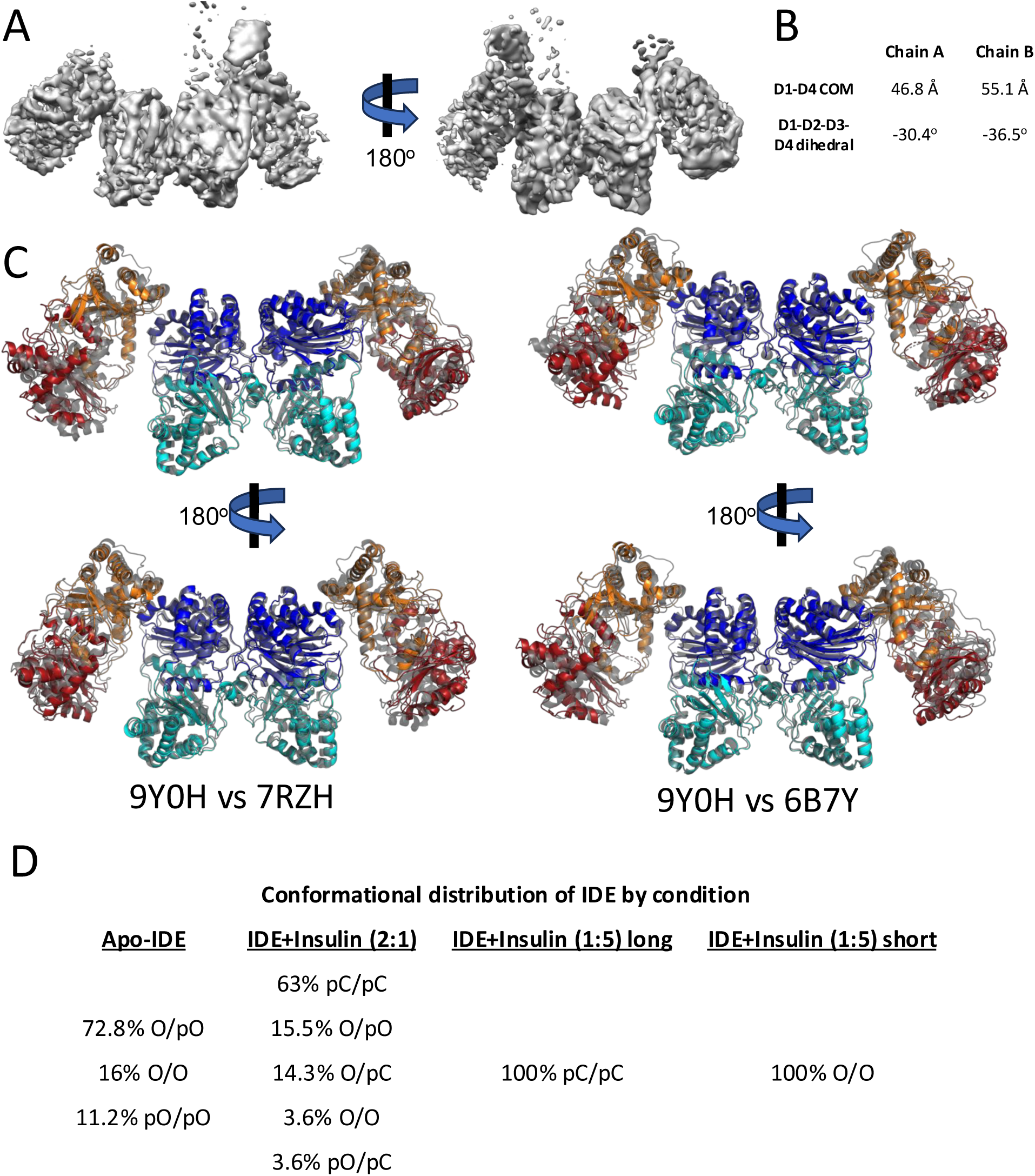
Time-resolved cryo-EM of IDE+insulin reveals a new O/O state. **(A)** 5.1 Å reconstruction of IDE+Fab rapidly mixed with a 5x molar excess of insulin and vitrified with a mix-to-freeze time of 123 milliseconds. **(B)** Measurements (as in Fig. 2) of the distance between the D1-D4 centers-of-mass and dihedral angle formed by the D1-D2-D3-D4 centers-of-mass as indicators of the “openness” of the structure depicted in (A). **(C)** Alignment of the O/O state from our rapid-mixing dataset (domains colored as in Fig. 1) compared to the previously solved O/O states (gray, as labeled). **(D)** Overview of the conformational states adopted by IDE in all 4 available cryo-EM datasets with corresponding percentage of particles mapping to each state (Zhang *et al*., 2018)(this work).

### Coarse-grained MD simulations reveal intermolecular cross-β sheet formation between insulin and IDE in all four IDE domains

As cryo-EM proved unsuitable to explore the mechanism of insulin unfolding by IDE, we turned to MD simulations. As stated above, there is a notable discrepancy in the time frame between the open/close transition and insulin cleavage rate. This suggests that unfolding requires multiple cycles of the open-closed transition and, thus it is not currently realistic to use all-atom MD simulations to probe the unfolding of insulin by IDE. We then used a coarse-grained MD package, Upside, to probe the interaction between IDE and insulin. Upside performs the dynamics simulation for just six atoms per residue with the sidechains being represented by directional-dependent beads whose packing probabilities have the lowest free energy while still including key structure details to compute realistic forces (Faruk *et al*, 2022; Faruk *et al*, 2023; Jumper *et al*, 2018a, b). This allows rapid equilibration and sampling of the energy landscape when compared with the conventional all-atom MD simulation, achieving ∼10^4^ acceleration for the calculation time. Consistent with all-atom MD simulations, Upside simulations revealed the wide range of motions, including grid and hinge motions found in our all-atom MD simulation (Figure 8 – figure supplement 1). We observed the partial unfolding of insulin in our Upside simulations yet, importantly, the unfolding did not disrupt the inter-or intra-molecular disulfide bonds (Figure 8 video 1).

**Figure 8:**
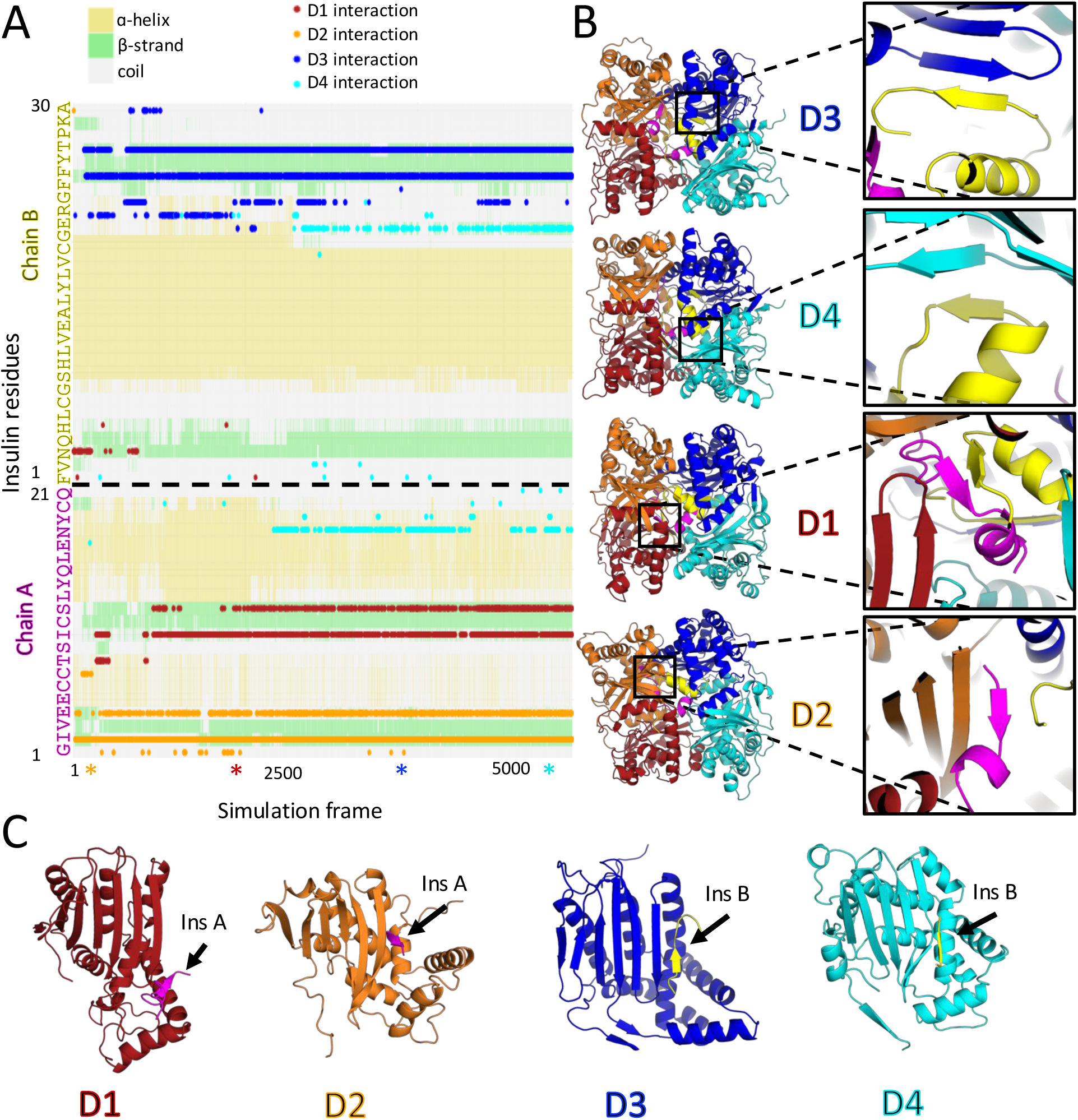
Upside analysis of IDE-insulin interactions. **(A)** Transient cross-β-strand interactions between insulin and IDE. Secondary structure (gray: coil, green: β-strand, yellow: α-helix) for each insulin residue (Y-axis) over the course (X-axis) of a representative Upside simulation. Points indicate predicted hydrogen bond interactions between IDE and insulin, colored by which domain of IDE that insulin residue is interacting with (D1: red, D2: orange, D3: blue, D4: cyan). Stars below the X-axis indicate frames from which structures were extracted for panel B. **(B)** Structure of IDE/insulin complex in Upside simulations at frames indicated in (A). IDE domains colored as in (A), insulin shown in magenta (chain A) and yellow (chain B). Inset highlights the intermolecular cross-β sheets formed between IDE and insulin within each of the four homologous IDE domains. **(C)** Individual domains of IDE, colored as in (A), extracted from (B) and oriented to demonstrate the structural conservation of intermolecular cross-β-sheet formation with bound insulin (colored as in B).

As insulin partially unfolds, it readily adopts β-strand structures—a key characteristic of amyloidogenic peptides—which can transiently form cross-β-sheets with the central β-sheet present in all four domains of IDE (Figure 8A-B). Specifically, we observed that cross-β-sheet interactions occur at distinct regions of insulin: the N-terminus of insulin chain A interacts with the IDE D2 domain near the substrate N-terminal anchoring site (also known as the exosite); the middle portion of insulin chain A interacts with the IDE D1 domain; and the C-terminal regions of insulin chain B interact with the D3 and D4 domains of IDE. While the interactions of insulin with IDE-C domain are novel, those with IDE D1/D2 domains are structurally similar to those previously observed (Guo *et al*, 2010; Manolopoulou *et al*., 2009; Ren *et al*, 2010; Shen *et al*., 2006). The labile and transient β-strand-rich interactions have been shown to mediate diverse protein-protein interactions, e.g., the assembly of intermediate filaments (Eibauer *et al*, 2024; Zhou *et al*, 2021).

This interaction provides molecular insight into how IDE recognizes and binds amyloidogenic peptides with diverse sequences (Figure 8C). IDE is capable of forming cross-β interactions with many regions of insulin, primarily mediated by backbone-to-backbone contacts that do not rely on specific amino acid sequences. The structural arrangement of the four conserved domains of IDE enables it to interact with multiple regions of its substrates, such as insulin, which are predisposed to forming β-strands (Figure 8C). We term this type of interaction “β-grabbing,” and propose that it is a key mechanism by which IDE unfolds insulin. This interaction effectively tethers the substrate to both the IDE-N and IDE-C domains, facilitating the interdomain motions of IDE that promote substrate unfolding.

## Discussion

Our results allow us to put forth a refined model describing how IDE recognizes amyloid peptides that are diverse in size and shape (Fig. 9). The catalytic cycle of IDE starts with at least one subunit of dimeric IDE adopting the open state, exposing the interior of the catalytic chamber to potential substrates (Zhang *et al*., 2018). The unconstrained motion of IDE-N likely facilitates substrate interaction in a variety of initial orientations. IDE-N and IDE-C have negatively and positively charged surfaces, respectively, and are able to attract peptide substrates with complementary charge profiles (Shen *et al*., 2006; Zhang *et al*., 2018). Such peptides have a high dipole moment and are often prone to aggregation. However, in the open state, the door subdomain, which contains the key catalytic residues, is highly flexible, rendering the open state catalytically incompetent (McCord *et al*., 2013; Zhang *et al*., 2018). Proper positioning and unfolding of substrate is necessary to stabilize the cleft prior to catalysis. Substrate binding has been suggested to enhance IDE closing through charge complementarity (Shen *et al*., 2006). It is likely that the rotation of IDE-N permits a range of closing pathways to accommodate non-optimal substrate binding geometry (Fig. 4). If the closing geometry permits R668 to interact with E381 or D309, the closing reaction continues to completion. If the closing geometry does not permit the R668-mediated interactions, this could be a signal of improper substrate binding, allowing IDE-N to open and either release improper substrate or reposition for another closing attempt. Once closed, IDE-N is capable of rotating against IDE-C mediated by charge-swapping events at the IDE-N/C interface (Fig. 6). The combined motions associated with the open to closed transition and rotation between IDE-N and IDE-C can lead to the distortion of the secondary structure of bound substrate. Furthermore, the transient interaction of insulin with the catalytic chamber of IDE such as β-grabbing via the β−strand pairing should further stress the integrity of bound substrate (Fig. 8). Together, these forces should facilitate unfolding and proper positioning of substrate for cleavage. Our data indicates that IDE is able to accommodate any possible orientation of insulin tethering within the catalytic site (Fig. 2C), suggesting that IDE is able to cut both chains of insulin in rapid succession without requiring significant repositioning (Manolopoulou *et al*., 2009). IDE then opens to release cleavage products and the cycle begins anew.

**Figure 9:**
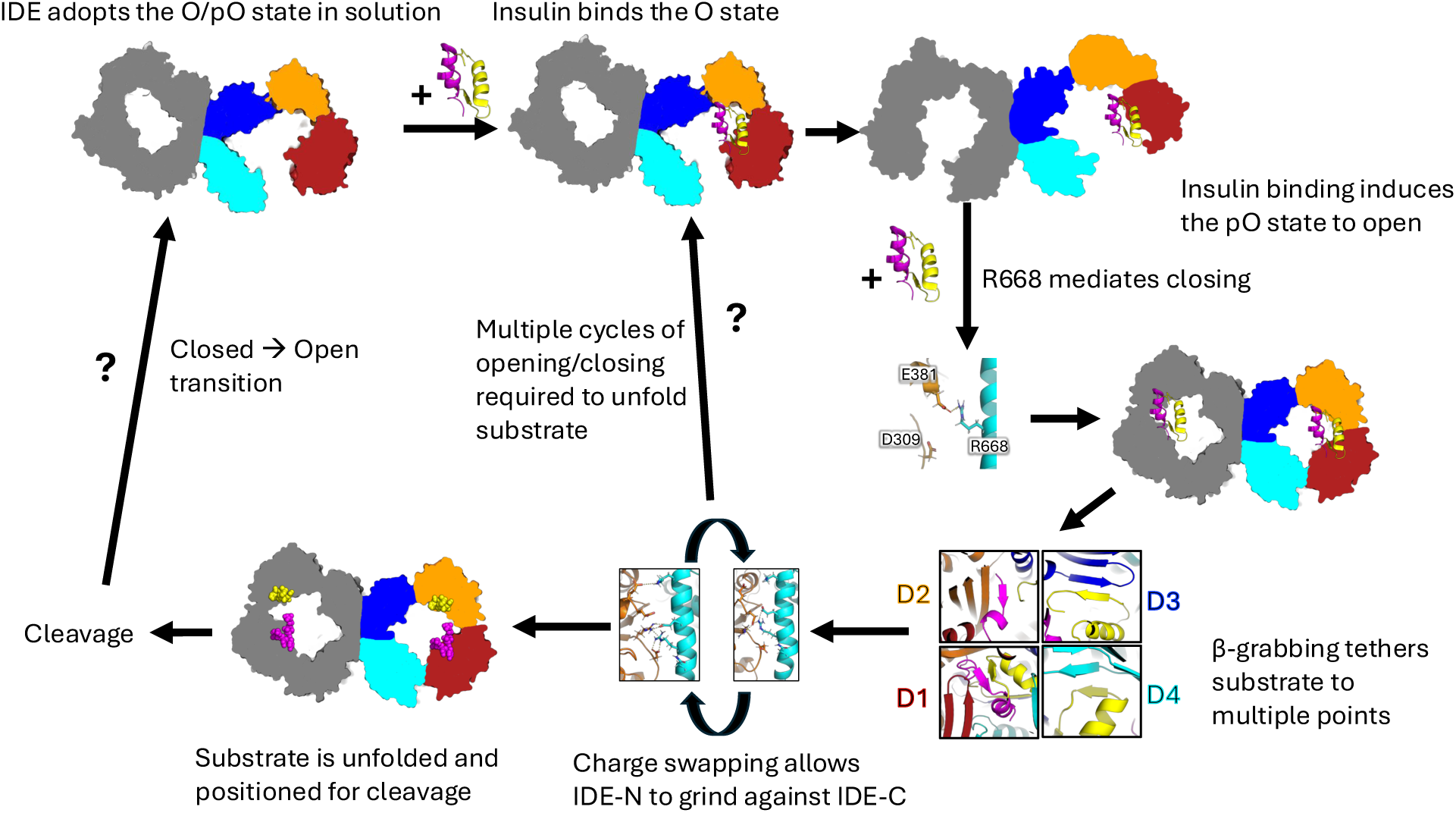
Model for the catalytic cycle of IDE. The details are described in discussion. For simplicity, only the scenario involving the addition of a second insulin molecule is illustrated, although this is not required for the catalytic cycle. The mechanism by which IDE transitions between closed and open states remains unknown and is therefore indicated with question marks.

IDE utilizes both unfoldase and protease activities to degrade clinically-relevant peptides, including three glucose-regulating hormones with contrary effects – insulin, amylin, and glucagon – and Aβ, the accumulation of which is associated with the progression of Alzheimer’s disease. This has made IDE an attractive target for therapeutic intervention (Maianti *et al*., 2019; Tang, 2016). One major challenge is how to better control selectivity when the substrates IDE degrades are highly diverse in sequence and structure. It remains unknown how IDE unfolds substrates prior to cleavage. Our studies characterize the conformational dynamics of IDE and offer insight into the catalytic mechanisms, revealing that IDE-N rotates against IDE-C. We envision that this motion works in concert with the open-close transition to reposition and unfold substrates prior to cleavage. This rotation appears to be largely unconstrained in the open state and may play a role in promoting substrate capture. In the closed state, we found the rotational motion to be mediated by charge-swapping events at the IDE-N/C interface. These events are supplemented by interactions between hydrophobic patches that provide a non-specific interaction interface and allow the domains to slide against one another. We found that IDE-N/C interactions were mediated by the key residue R668, and mutation of this residue altered the enzymatic activity and conformation of IDE, likely by impairing the open-close transition. Our data suggests that, while the R668A mutation globally impaired IDE activity, R668A exhibits significantly different kinetic profiles towards well-folded substrates (insulin) compared to unstructured substrates (Aβ). This highlights the complexity of the IDE catalytic cycle as we envision that IDE conformational dynamics can regulate solvent access to the catalytic chamber and substrate binding, unfolding, and cleavage are all intimately linked.

Our work adds to the expanding body of literature that integrates computational and experimental approaches (Arenz *et al*, 2016; Hirschi *et al*, 2021; Jiang & Doudna, 2017; Palermo *et al*, 2016; Roh *et al*, 2020; Saha *et al*, 2022). By employing multibody analysis, we identified the dominant sources of structural heterogeneity within our cryo-EM particle population, uncovering that the conformational dynamics of IDE involve a substantial rotational component that is not evident from ensemble structure analysis alone. We showed that all-atom MD simulations can address limitations of cryo-EM single particle analysis when investigating the role of residue R668. Because R668 forms a network of polar and electrostatic interactions rather than exclusive partnerships, dynamic hydrogen bonding networks at the IDE-N/C interface—particularly those involving R668—contribute to conformational heterogeneity, which further restricts the achievable resolution in cryo-EM analysis. Furthermore, vitrification by freezing proteins in liquid ethane can significantly alter the entropic contribution of R668 through conformation-dependent promiscuous contacts. By combining cryo-EM and MD simulations, we gained insight into how flexible interactions between IDE-N and IDE-C facilitate domain movements and highlighted the critical role of R668 in mediating these interactions. We also used MD simulations to demonstrate altered open-closed dynamics in the R668A mutant compared to the wild-type enzyme and validated these findings with experimental enzymatic kinetic assays and SEC-SAXS experiments.

At its core, cryo-EM is an ensemble method of structure determination, averaging tens of thousands of particles together to generate consensus structures. While methodologies have been well-established to separate out discrete conformational classes, proteins that exhibit constant gradients of structural heterogeneity remain problematic, although significant attention has been devoted to the issue in recent years, with varying levels of success (Nakane *et al*., 2018; Punjani & Fleet, 2021; Zhong *et al*, 2021). Currently, the best approaches use various dimensionality reduction techniques to analyze the principal components of particle heterogeneity. While a substantial innovation, it remains to be seen how the principal components of structural heterogeneity, derived from a series of static snapshots of a particle at the time of freezing, correlate with the real molecular motions of a protein in an aqueous environment. For many systems, especially enzymes of clinical significance, proper characterization of the conformational dynamics and transitional states of the protein are necessary for a full understanding of protein function and lays the foundation for future therapeutic development. MD simulations can provide this information, yet such studies are often met with skepticism by experimental researchers. By integrating MD simulations with experimental techniques, the weaknesses of one technique can be compensated for by the strengths of other techniques to provide comprehensive information about the system of interest. While this integrative approach has proven beneficial across a wide range of systems, including proteorhodopsin, the ribosome, and rotary ATPases, perhaps the best example of this approach is the CRISPR-Cas9 system, where an exhaustive number of studies have combined cryo-EM, crystallography, and traditional biophysical techniques with traditional and accelerated MD methods to drive the field forward at an astounding pace (Arenz *et al*., 2016; Hirschi *et al*., 2021; Jiang & Doudna, 2017; Palermo *et al*., 2016; Roh *et al*., 2020; Saha *et al*., 2022).

Our integrative structural analyses provide the structural basis for IDE conformational dynamics and their roles in IDE catalytic cycle and the framework for future studies. We speculate that multi-cycles of open-close transition and rotation driven by IDE conformational dynamics will be required for its unfoldase activity. However, the kinetic timescale of this reaction is fast, estimated to be ∼500 milliseconds rendering investigation challenging experimentally while the timescale is long for all-atom MD simulation (Manolopoulou *et al*., 2009). Our cryo-EM analyses also shed light on IDE allostery (McCord *et al*., 2013; Song *et al*., 2011; Zhang *et al*., 2018) (Fig. 9). The IDE dimer primarily adopts the O/pO conformational state (Zhang *et al*., 2018). Through time-resolved cryo-EM analysis, our data suggest that, upon substrate binding to the O state, the substrate-inaccessible pO state undergoes allosteric opening, allowing this subunit to capture its substrate. Additionally, the prolonged incubation of IDE with sub-stoichiometric amounts of insulin results in the predominance of the pC/pC state, suggestive of coordination in substrate capture and open-closed transition. Collectively, these positive allosteric transitions would enhance the capacity of IDE to capture and degrade its substrates more efficiently, though the underlying mechanisms remain unclear. Future integrative structural studies are required to further elucidate and control the IDE catalytic cycle, enabling substrate-specific modulation of IDE activity and advancing its therapeutic potential.

## Materials and Methods

### Expression and purification of human IDE

Cysteine-free IDE (referred to as IDE in this study) was expressed in *E. coli* BL21 (DE3) cells (at 25 °C for 20 h, 0.5mM IPTG induction using T7 medium). The R668A mutation was introduced into this construct by site-directed mutagenesis with the following primer pair: CCGGAAATTGTTAAGAGATGCCATATATGCTTCTTTG (forward) CAAAGAAGCATATATGGCATCTCTTAACAACCGGGC (reverse) and verified by sequencing. Recombinant IDE proteins were purified by Ni-NTA, source-Q, and Superdex 200 columns as previously described (Zhang *et al*., 2018). Protein was aliquoted and stored at −80° C.

### Cryo-EM data collection and analysis

Thawed IDE aliquots were further purified by Superdex 200 chromatography using buffer containing 20 mM HEPES, pH 7.2, 150 mM NaCl, 10mM EDTA and then mixed with Fab_H11-E_ at an equal molar ratio. Fab_H11-E_-IDE complex was purified by Superdex 200 chromatography in the presence of insulin at molar ratio IDE:insulin=2:1. Insulin was purchased from SIGMA (91077C). All grids, 300 mesh carbon or gold holey nanowire grids were plasma cleaned with O_2_ and H_2_ for 10 secs using a Solarus plasma cleaner (Gatan) and plunged at 133 milli-second using Chameleon, Spotiton-based technology. All images were acquired using a Titan Krios microscope (FEI) operated at 300KeV with a Gatan K3 direct electron detector (Gatan) in counting mode. Images were automatically acquired using Leginon (Suloway *et al*, 2005). Images were processed using software integrated into RELION3 (Zivanov *et al*, 2018). Frames were aligned using MotionCor2 (Zheng *et al*, 2017) software with dose weighting, CTF was estimated using Gctf (Zhang, 2016), particles were picked and extracted automatically using RELION. Particle stacks were processed through several rounds of 2D and 3D classification. Selected classes were then processed for high-resolution 3D refinement. The flowchart and detailed data processing is summarized in Figure 2 figure supplement 1. Finally, the overall map was improved by particle polishing in RELION and sharpening. The final resolution was estimated using Fourier Shell Correlation (FSC=0.143, gold-standard). The density fitting and structure refinement was done using UCSF CHIMERA (Pettersen *et al*, 2004), COOT (Emsley & Cowtan, 2004), REFMAC5 (Murshudov *et al*, 2011) and PHENIX (Adams *et al*, 2010). Time-resolved cryo-EM grids by rapidly mixing IDE+Fab with insulin were prepared using the Spotiton device as previously described (Budell *et al*, 2021; Dandey *et al*., 2020). In brief, IDE+fab and insulin were thawed on ice and diluted in 20 mM HEPES, pH 7.2, 150 mM NaCl, 10 mM EDTA to 25 µM and 127.5 µM, respectively. Self-wicking, nanowire grids were plasma treated (Gatan Solarus) at 5 W in hydrogen and oxygen for 1-2 minutes (Wei *et al*, 2018). Each sample was aspirated into separate piezodispensers and applied sequentially in 70-100% relative humidity at 22-25C onto plunging grids just before cryogen entry. The maximum constant velocity of the plunging robot was adjusted to obtain the target sample mixing times of 123 milliseconds. The data was collected and processed as above, in Relion5, with Blush regularization being employed during 3D refinement.

### Multibody analysis

We defined three rigid bodies as shown in figure 2A for the analysis. The size of the user-defined bodies has been theorized to play a significant role in the success of multibody refinement, most likely due to the fact that larger bodies will have a stronger signal-to-noise ratio (Nakane *et al*., 2018). With a Fab bound to each of IDE-N within IDE dimer, we assessed the impact of body size on multibody refinement by examining how density corresponding to 2 F_v_ regions, 1 F_v_ region, or no F_v_ regions would affect the multibody refinement. Fab has a variable region (F_v_), and a constant region (F_c_) and we chose to subtract and mask out the density corresponding to the Fab F_c_, as we speculated that any motion between the Fab F_v_ and F_c_ domains would bias the results of the multibody refinement. The quality of the multibody results was assessed based on the subsequent improvement in the Coulomb potential map quality and calculated resolution. In the absence of any F_v_ density, the maps resulting from multibody refinement had the best calculated resolution, yet the map quality was quite poor overall; much of the density appears globular and featureless, particularly in the IDE-N regions (Figure 2 – figure supplement 4). Conversely, when F_v_ density was present on both IDE-N bodies, the resolution of the multibody output maps (4.5 Å) was worse than the resolution of the input map (4.3 Å). Thus, we found that the greatest improvement occurred when the F_v_ density was present on only the exterior body of the pO subunit. In this case, multibody refinement improved both the calculated resolution and density quality.

Following multibody refinement, the structural heterogeneity for each body was analyzed along six principal component vectors, resulting in 18 vectors describing the primary components of structural variation within the IDE dimer. We analyzed the variation along these vectors as a proxy for protein motion. The results of the multibody analysis indicate the degree of structural variation that is explained by motion along each of the primary component vectors. For all IDE states, we observed no single component vector could explain the majority of structural variance (Supplementary files 3-8). Given the underlying assumptions of multibody refinement, there is a question of whether any observed motions are biologically relevant or simply the result of improper particle alignment. We reasoned that the greater the degree of variation explained by motion along a specified component vector, the greater the likelihood was that the motion would be biologically relevant. Therefore, we focus on the top 9 component vectors, which cover ∼75% of total variance for each state, in greater detail.

### Molecular dynamics simulations

All-atom MD simulations were prepared from the O/pO structure of IDE (PDB: 7RZK) and necessary CHARMM force field PARAM36 files using QwikMD (Ribeiro *et al*, 2016). Gaps in the structure were modeled in using the AlphaFold model of IDE. A three-step minimization-annealing-equilibration process was used to generate an equilibrated system. Simulations were performed under NPT (constant numbers of particles N, pressure P, and temperature T) conditions at 310 K and 1 atm with periodic boundary conditions in NAMD3.0 (Phillips *et al*, 2005). Explicit solvent was described with the TIP3P model (Jorgensen *et al*, 1983). Custom TCL scripts in VMD were used to calculate the D1-D4 centers-of-mass distance and D1-D2-D3-D4 centers-of-mass dihedral angle (Supplementary file 9). Residues 964-988 in D4 that are mostly absence in all static structures of IDE were built using Alpha-fold, which is highly flexible in our MD simulations. They were omitted from the analysis of MD simulations, as the inclusion of this flexible loop significantly altered the center-of-mass independent of global conformational change.

Initial structure of insulin bound IDE, for Upside simulations, was generated using PYMOL that started with IDE closed monomer (PDB code: 6BF8). Intact insulin (PDB code: 4INS) was fixed by mutation of residue 30 of insulin chain B back to threonine and modeled into the catalytic chamber based on the crystal structure of IDE in complex with the partially unfolded insulin (PDB code: 2WBY). The upside simulation was run with 28 replicates at T_Upside_=0.8 using Upside 1.0 force field parameters and our total upside simulation time is approximately 28 μsec (Faruk *et al*., 2022; Faruk *et al*., 2023; Jumper *et al*., 2018a, b). We estimated the time step of Upside to be ∼0.1 picosecond based on folding rates and chain motions. Custom R, and Python scripts were used for data analysis and presentation (Supplementary files 10-12).

### Enzymatic assays

IDE constructs were exchanged into activity buffer comprised of 25 mM Tris-HCl pH 7.5, 150 mM NaCl, and 10 mM ZnCl_2_. MCA-RPPGFSAFK(Dnp) was purchased from Enzo Life Sciences (BMLP2270001), resuspended in dimethyl sulfoxide at a concentration of 5 mM and diluted to 5 μM with activity buffer. For activity measurements, 5 μM MCA-RPPGFSAFK(Dnp) was mixed with the desired IDE construct at concentrations ranging from 1-100 nM in the presence or absence of 1 mM ATP in a 200 μl reaction. Fluorescence was monitored every 30 sec for 30 minutes with excitation/emission wavelengths of 320/405 nm at 37° C. Initial velocity was calculated during the linear range. Insulin was purchased from MP Biomedicals (#0219390010), resuspended in 0.01N HCl at 1 mM and diluted to desired concentration with activity buffer. For competition assays, 5 nM IDE construct and 5 μM MCA-RPPGFSAFK(Dnp) was mixed with insulin ranging from 0-100 μM. Fluorescence was monitored every 30 sec for 30 minutes with excitation/emission wavelengths of 320/405 nm at 37° C. Initial velocity was calculated during the linear range for each concentration of insulin and normalized relative to the velocity of the respective construct in the absence of insulin to maintain consistency across plates, yielding values for relative inhibition which were plotted vs [insulin] or [Aβ] and fit to the Michaelis-Menten equation in xmgrace to generate apparent K_i_ values. All experiments were performed in triplicate.

### Size-exclusion chromatography coupled small-angle X-ray scattering

SAXS/WAXS data collection was employed the Life Sciences X-ray Scattering (LIX) Beamline at the National Synchrotron Light Source II (NSLS II) at Brookhaven National Laboratory in Upton, NY. Briefly, 60ul of samples in solution were pipetted into PCR tubes, placed into a Bio-Inert Agilent 1260 Infinity II HPLC multisampler and measured using the isocratic SEC-SAXS format at the beamline (Yang *et al*, 2021). 50ul of sample was injected into the Phenomenex Biozen dSEC2 3 μm bead size with 200Å pore size column was utilized at a flow rate of 0.35mL/min for 25 minutes. SAXS and WAXS data are collected simultaneously on a Pilatus 1M (SAXS) and Pilatus 900K (WAXS) detectors with a 2 second exposure (Yang *et al*, 2020). Data from both detectors is then scaled and merged. Intensity is normalized using the water peak height at 2.0 Å ^-1^. Data processing and analysis was performed using py4xs and lixtools in jupyter lab. Buffer frames 100-125 were averaged and used for subtraction of averaged frames under the peak of interest.

## Data Availability

Cryo-EM maps and refined models have been deposited to the EMDB and PDB, respectively with the following accession numbers:

O/O state: EMDB-24760, PDB 7RZH

O/pO state: EMDB-24759, PDB 7RZG

pO/pC state: EMDB-24757, PDB 7RZE

O/pC state: EMDB-24758, PDB 7RZF

pC/pC state: EMDB-24761, PDB 7RZI

Time-resolved O/O state: EMDB-72393, PDB 9Y0H

Cryo-EM micrographs have been deposited to the Electron Microscopy Public Image Archive (EMPIAR) under the following accession numbers:

IDE+insulin in a 2:1 ratio: EMPIAR-11610

Time-resolved IDE+insulin: EMPIAR-12968.

## Acknowledgement

We would like to thank our colleagues at the University of Chicago: Adam Antoszewski and Aaron Dinner for providing us with the data from their insulin unfolding simulations, as well as Rong Shen for general guidance with the MD simulations, and Jessica Martinez Martinez for assistance with cloning the IDE R668A construct. We would also like to thank James Byrnes at the LiX beamline at Brookhaven National Laboratory for collecting SEC-SAXS data. Computer resources came, in part, from an allocation on the Beagle3 computing cluster at the University of Chicago. This work was supported by the NIH grant GM121964 to W.-J. Tang. The Simons Electron Microscopy Center is supported by a grant from the Simons Foundation (SF349247). Use of the Advanced Photon Source was supported by the U.S. Department of Energy, Office of Basic Energy Sciences, under contract No. DE-AC02-06CH11357. The LiX beamline is supported by NIH grants P30GM133893 & S10 OD012331, DOE grants KP1605010 & DE-SC0012704.

