## Supplementary figures and files for "Characterization and modulation of human insulin degrading enzyme conformational dynamics to control enzyme activity"

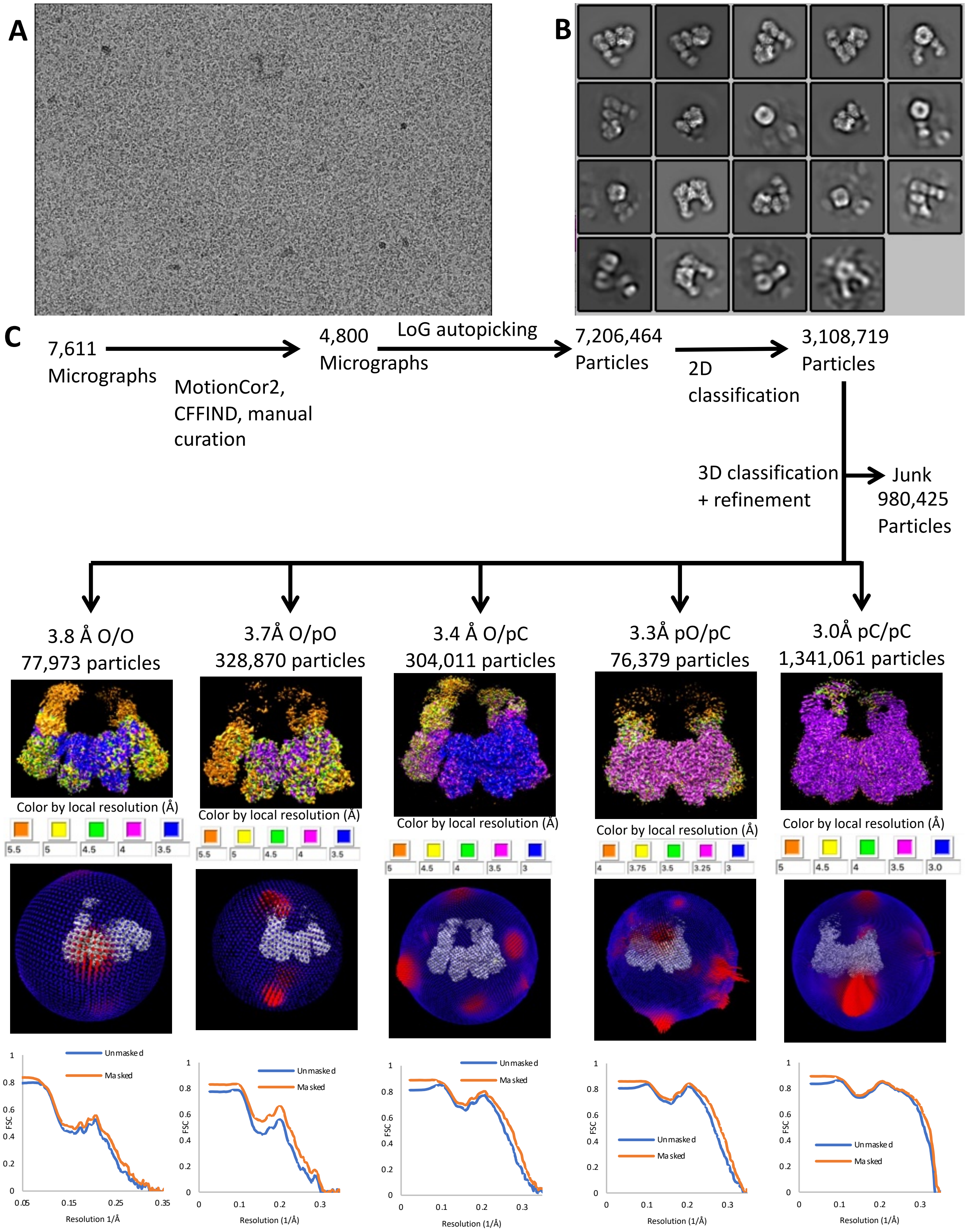


Figure 2 - figure supplement 1 – Cryo-EM summary. (A) Representative micrograph. (B) selected 2D classes. **(C)** processing workflow, structures colored by local resolution, angular distribution, and FSC curves.

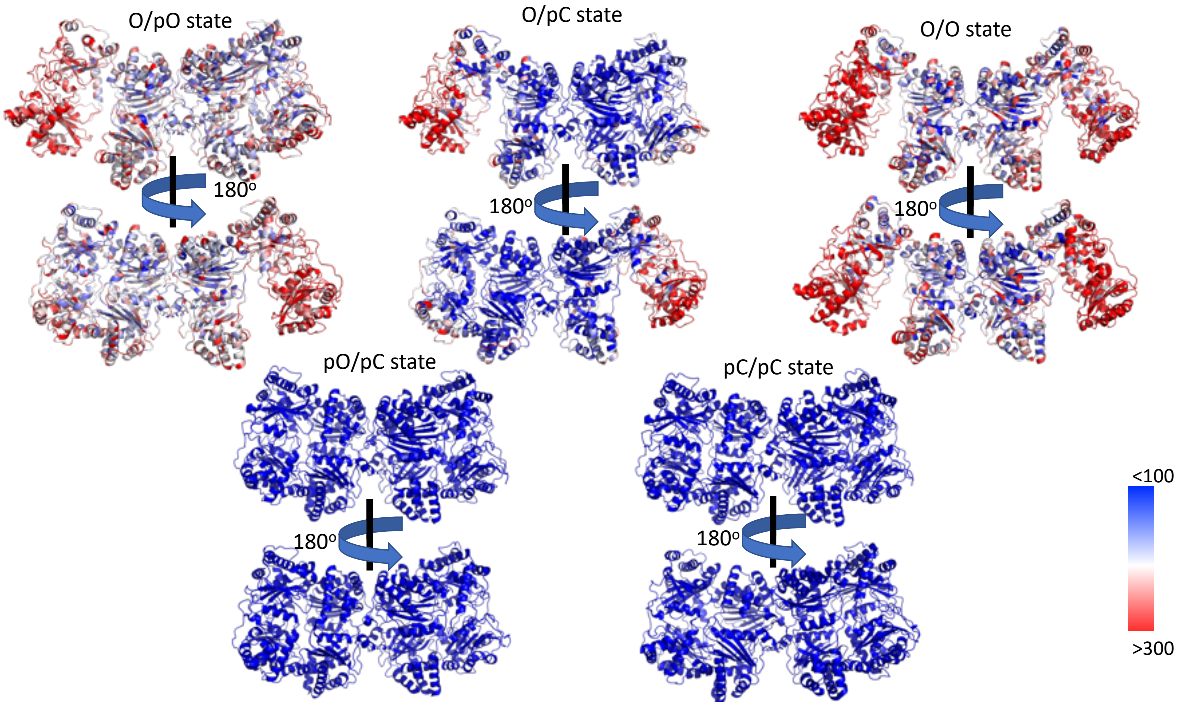


**Figure 2 - figure supplement 2 – B-factor analysis of 2:1 IDE:insulin complex cryo-EM structures.** Colored blue-white-red to show B-factor distribution from <100 to >300. The O state, particularly the D1 domain consistently demonstrates higher B-factors than the rest of the structure.

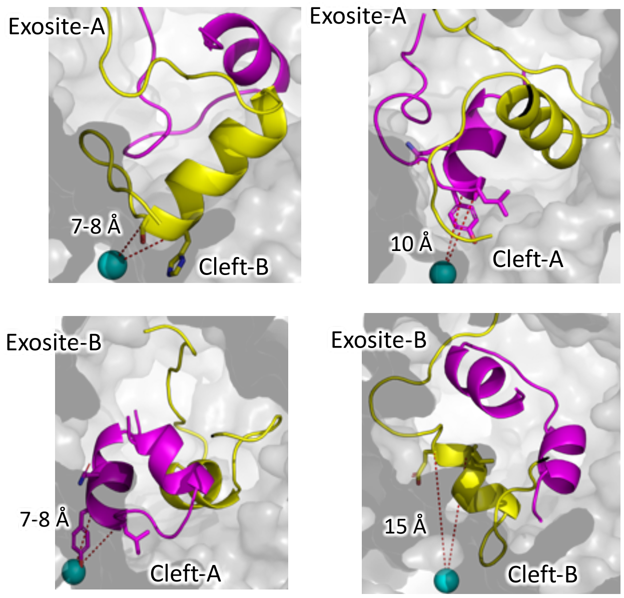

**Figure 2 - figure supplement 3 – Partially unfolded structures of insulin docked into the closed chamber of IDE.** Insulin colored by chain (A magenta, B yellow), catalytic zinc shown as cyan sphere. Distance from catalytic zinc to cleavage sites shown as red dashed line. Distance to nearest cleavage site listed. IDE shown as gray surface.

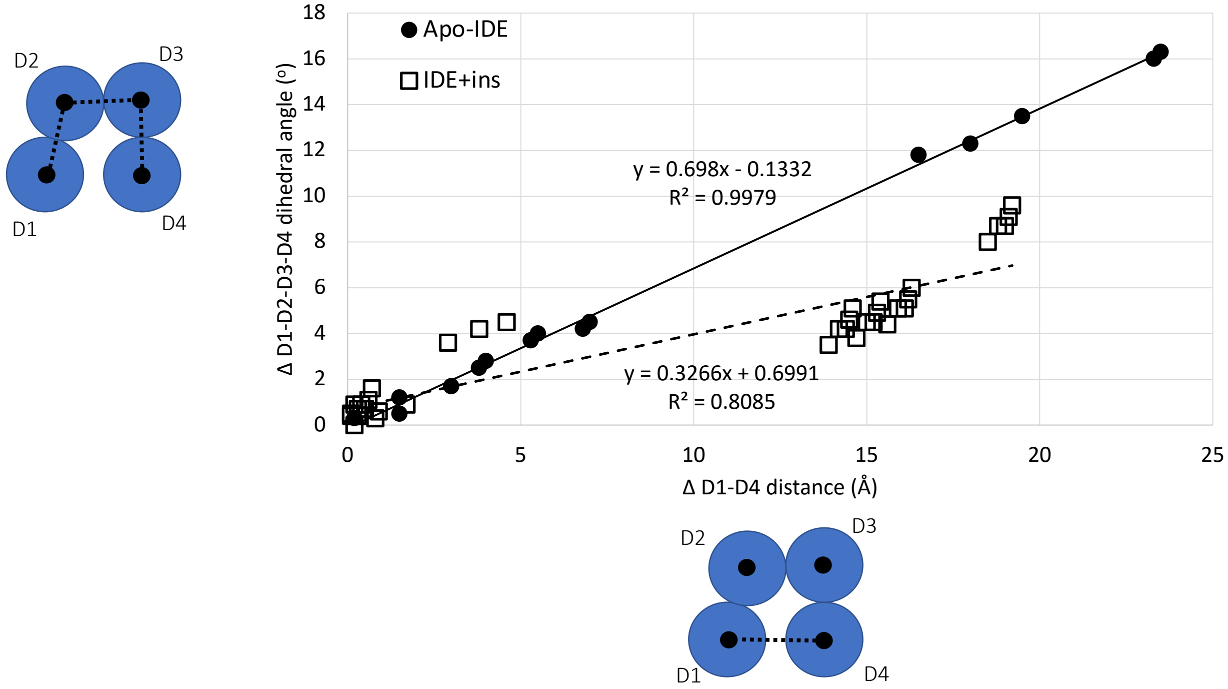


**Figure 3 - figure supplement 1 – Conformational space of ensemble structures.** The absolute value for the change in D1-D4 distance, and D1-D2-D3-D4 dihedral angle between available apo or insulin-bound structures of IDE in the Protein Databank was plotted to understand a potential open-close transition pathway. Structures with bound inhibitors or non-insulin substrates were omitted. For any given structure, the D1-D4 distance and D1-D2-D3-D4 dihedral angle were calculated from the domain centers of mass, as shown in the schematic along each axis, in PyMOL. To generate the ΔD1-D4 distance, and ΔD1-D2-D3-D4 dihedral angle, the respective value was subtracted from each of the same values generated for every other structure within the group of analyzed structures and plotted as absolute values. This method produced a linear fit of >0.99 for the structures solved in the absence of insulin (black dots, solid black line), but a linear fit of only 0.81 for the structures solved in the presence of insulin (open squares, dashed line), suggesting that the presence of insulin stimulates a larger change in the D1-D2-D3-D4 dihedral angle.

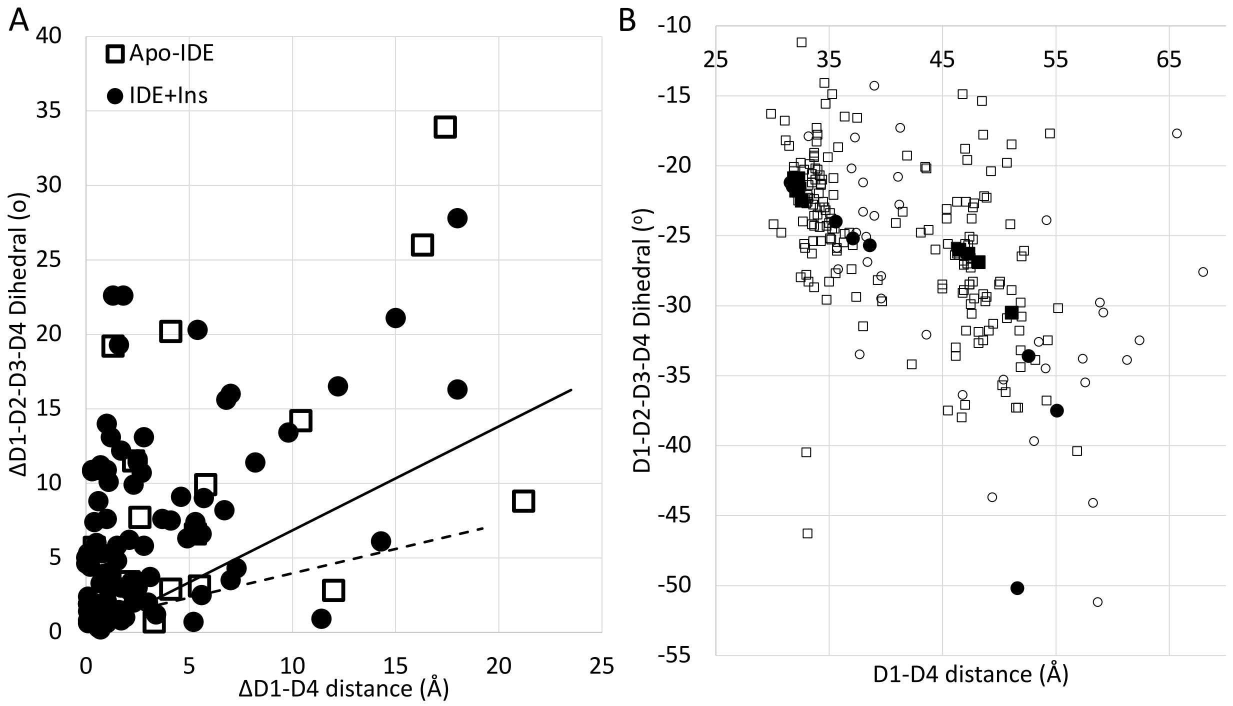


**Figure 3 - figure supplement 2 – Multibody analysis (A)** Open-close transition pathway predicted from multibody analysis. The solid and dash lines match those in Figure 3 – figure supplement 1, which depicts transition pathways derived from the static structures of apo-IDE and IDE+insulin. IDE domains were fit, as rigid bodies, into density maps representing the outermost limits of structural heterogeneity along the top 9 eigenvectors produced during multibody refinement, treated as static structures, and analyzed as in Figure 3 – figure supplement 1. Apo-IDE structures are shown as white squares, IDE structures in the presence of insulin are shown as black dots. Distribution of measurements from multibody analysis does not support the expected linear transition derived from analysis of the static structures, suggested that IDE exhibits an unexpectedly greater degree of rotational motion, as measured by the change in D1-D2-D3-D4 dihedral angle. **(B)** Conformational space sampled by top eigenvectors derived from Multibody analysis. Plot of the D1-D4 distance vs D1-D2-D3-D4 dihedral angle for the IDE cryo-EM structures. Values for the static structures of apo-IDE are shown as black dots, values for the static structures of IDE in the presence of insulin are shown as black squares. Each of the static structures was subjected to multibody analysis (see Methods). The resulting structures representing the outermost limits of structural variation along the top 9 eigenvectors for each structure were further analyzed and their D1-D4 distance and D1-D2-D3-D4 dihedral angle were plotted to yield the range of conformational space sampled by our cryo-EM particle population. Measurements derived from the multibody analysis of the apo-IDE structures are shown as white dots, measurements derived from the IDE+insulin structures are shown as white squares.

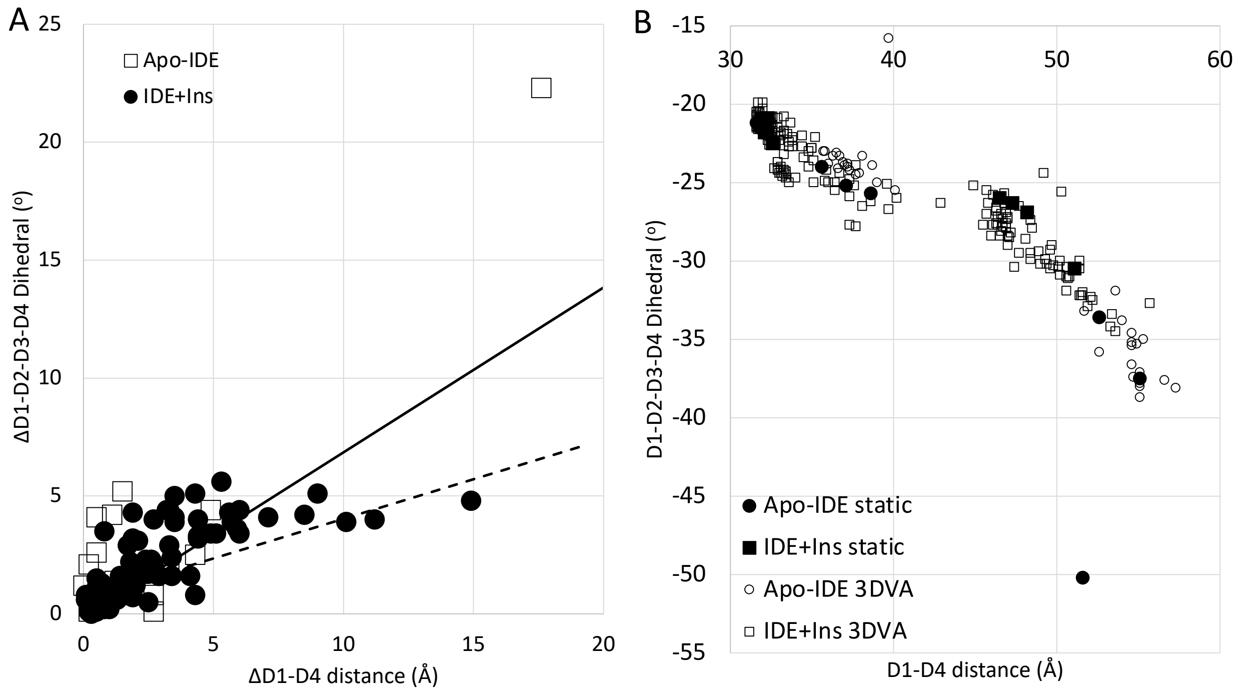


**Figure 3 - figure supplement 3 – Cryosparc 3D variability analysis (3DVA)_(A)** Open-close transition pathway predicted from 3DVA. The solid and dash lines match those in Figure 3 – figure supplement 1, which depicts transition pathways derived from the static structures of apo-IDE and IDE+insulin. IDE domains were fit, as rigid bodies, into density maps representing the outermost limits of structural heterogeneity along the top 9 eigenvectors produced during 3DVA, treated as static structures, and analyzed as in Figure 3 – figure supplement 1. Apo-IDE structures are shown as white squares, IDE structures in the presence of insulin are shown as black dots. The expected transition pathways derived from the static structures of apo-IDE and IDE+insulin are shown as a solid black line, and dashed black line, respectively. Distribution of measurements from 3DVA supports the observation from multibody analysis that IDE exhibits an unexpectedly greater degree of rotational motion, as measured by the change in D1-D2-D3-D4 dihedral angle. **(B)** Conformational space sampled by top eigenvectors derived from 3DVA. Plot of the D1-D4 distance vs D1-D2-D3-D4 dihedral angle for the IDE cryo-EM structures. Values for the static structures of apo-IDE are shown as black dots, values for the static structures of IDE in the presence of insulin are shown as black squares. Each of the static structures was subjected to identical 3DVA. The resulting structures representing the outermost limits of structural variation along the top 9 eigenvectors for each structure were further analyzed and their D1-D4 distance and D1-D2-D3-D4 dihedral angle were plotted to yield the range of conformational space sampled by our cryo-EM particle population. Measurements derived from 3DVA of the apo-IDE structures are shown as white dots, measurements derived from the IDE+insulin structures are shown as white squares.

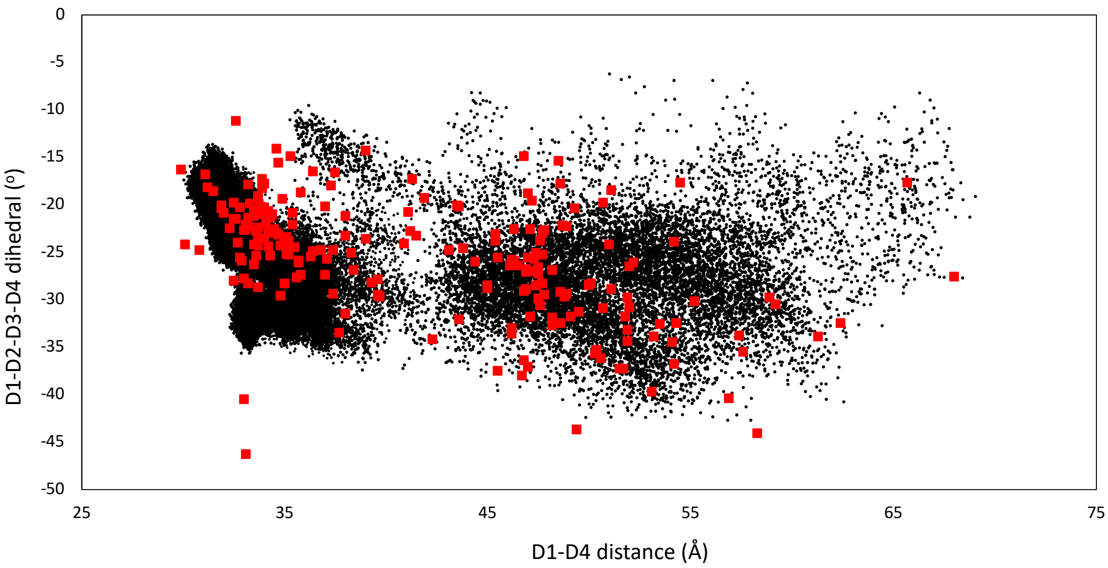


**Figure 4 - figure supplement 1 – Conformational space sampled by multibody analysis correlates well with MD simulations.** Plot of the D1-D4 distance vs D1-D2-D3-D4 to map the conformational space sampled by IDE. The extreme conformations along each of the top 9 eigenvectors for our multibody analysis (see Figure 3 – figure supplement 2), representing the conformational space sampled by our experimental particles, are shown as red squares. Black dots indicate the measurements of our WT IDE structure taken every 0.1 ns from our all-atom MD simulations, showing good agreement between the conformational space sampled experimentally and computationally.

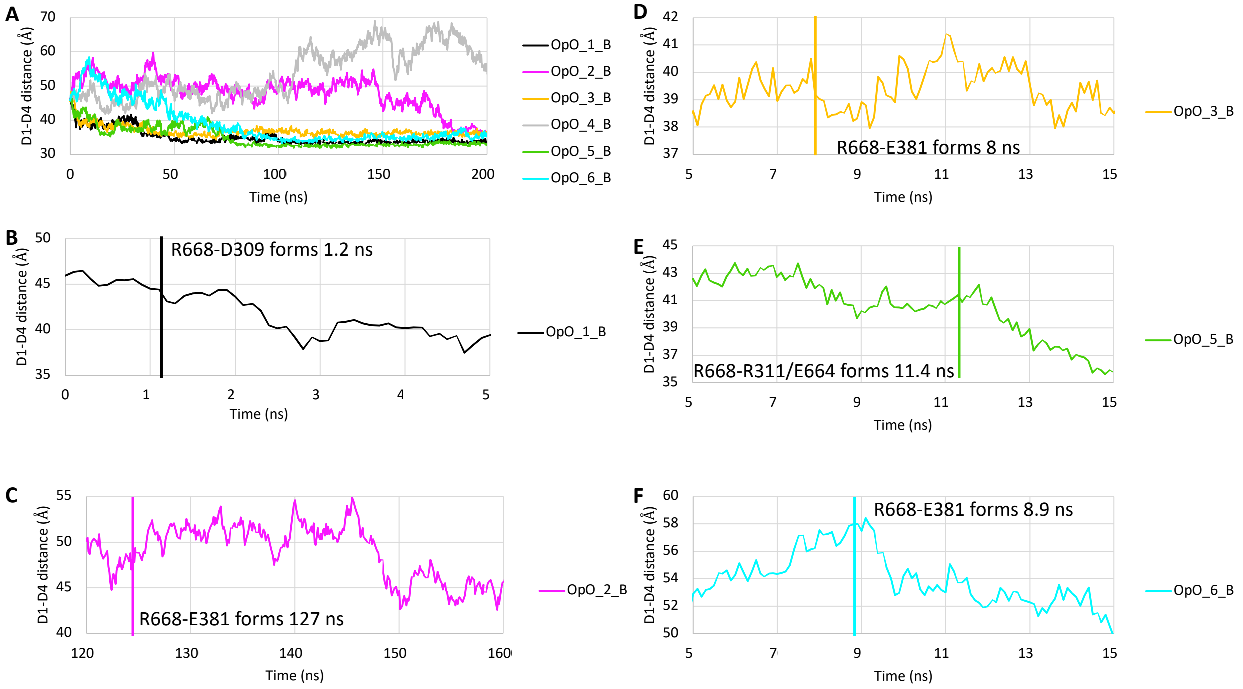


**Figure 4 - figure supplement 2 – R668-D309/E381 interactions are associated with closing. (A)** D1-D4 distance over the first 200 ns of our WT IDE MD simulations. **(B-F)** Changes in the D1-D4 distance over a shortened time window for the simulations where the open subunit closed. Line colors correspond to those in **(A)**. Vertical line indicates the timepoint where R668 first forms a lasting interaction with IDE-N (residue and time as indicated). In most simulations, this interaction precedes a rapid decrease in D1-D4 distance.

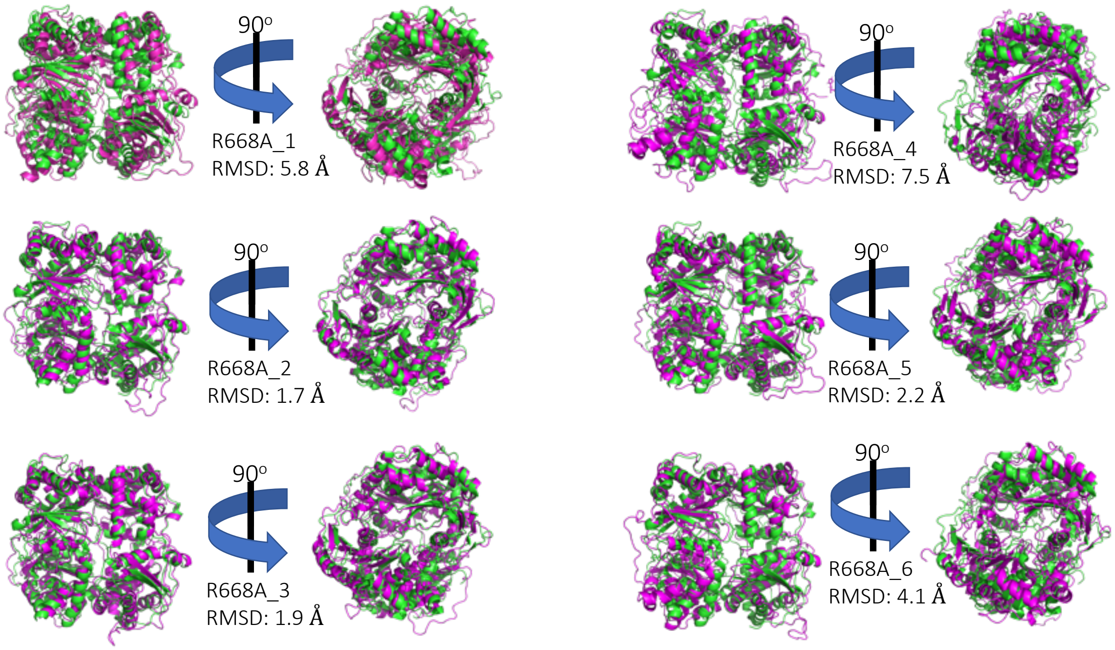


**Figure 4 – figure supplement 3 – Comparison of R668A all-atom MD O subunit end states compared**

**to ensemble closed state.** End state of the open subunit for each of the 6 IDE R668A MD simulations (magenta) aligned to the cryo-EM partial closed state structure (green, PDB: 7RZI) showing the altered closing geometry displayed by the mutant construct. RMSD for mainchain atoms as shown.

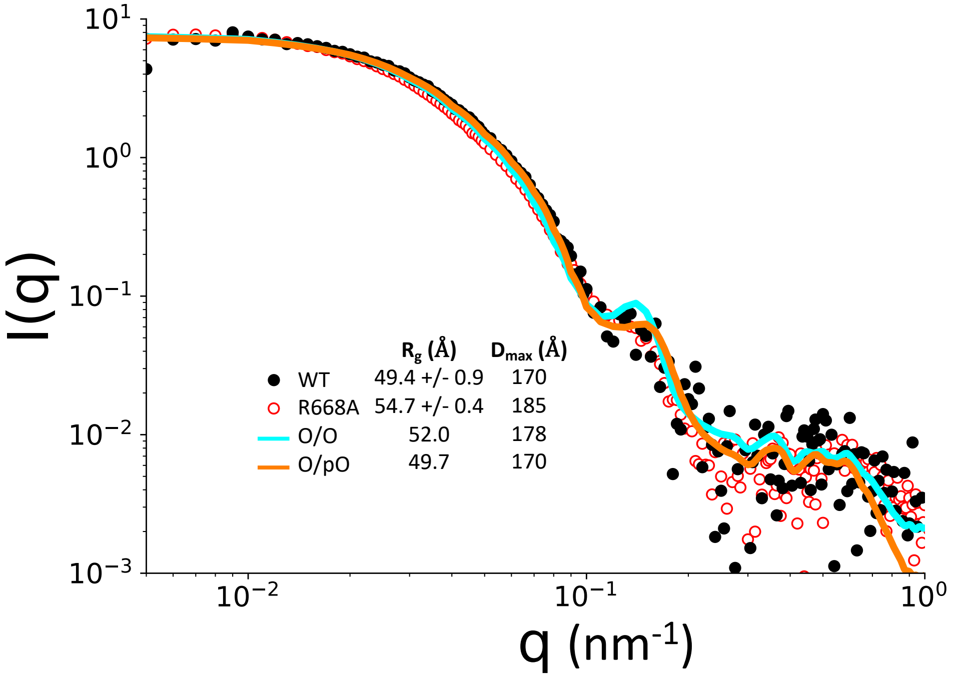


**Figure 5 – supplement 1 – Model fit to SEC-SAXS data.** SEC-SAXS data for WT IDE (black circles) and IDE R668A (open red circles) as shown in Fig. 4D. Calculated scattering patterns for experimentally determined structures were fit to the SEC-SAXS scattering profiles using CRYSOL.

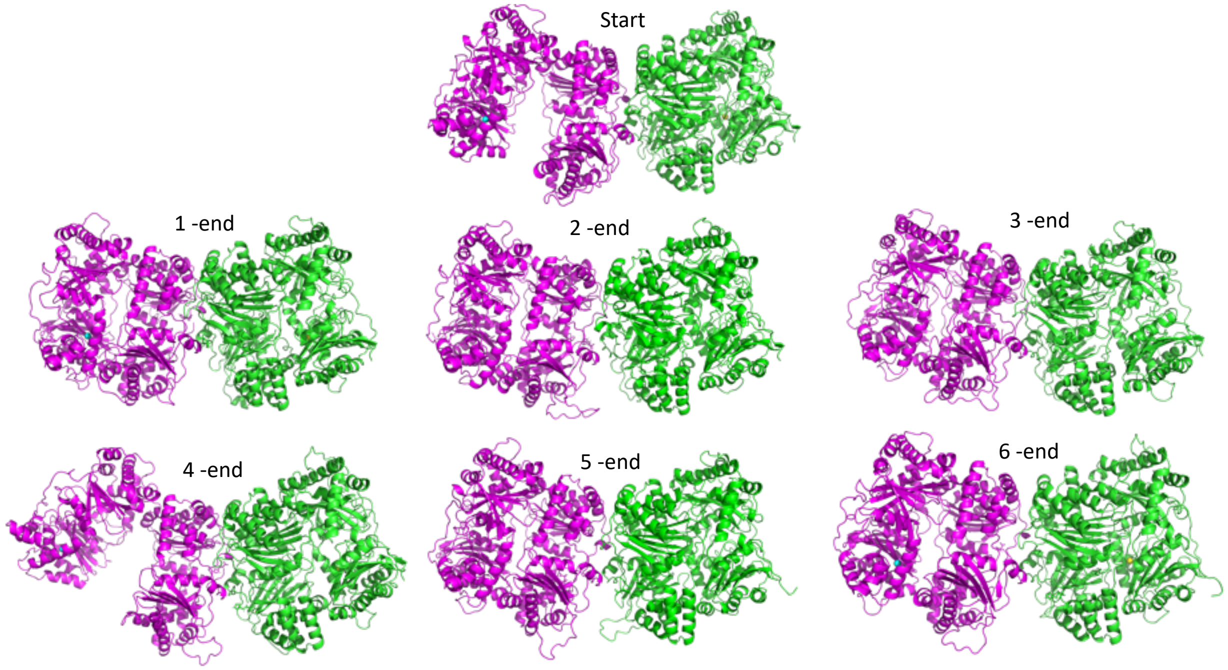


**Figure 6 – figure supplement 1 – Structures of WT IDE at the end of MD simulations.** The open subunit (magenta) closed in most simulations, but the closed subunit (green) did not open, leading to a nearly symmetric end state in 5 of the 6 simulations.

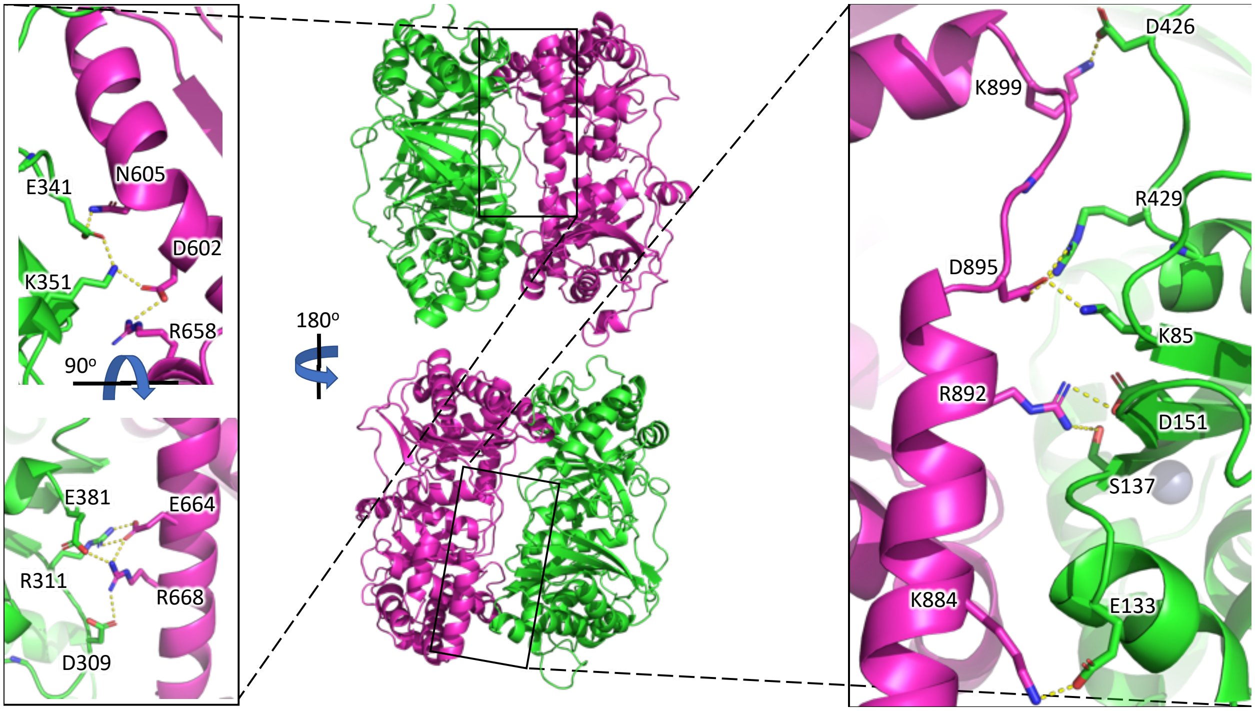


**Figure 6 – figure supplement 2 – IDE-N/C interface networks derived from all-atom MD simulations.** Key residues forming a dynamic hydrogen bonding network between IDE-N (green) and IDE-C (magenta). Left inset shows key D2-D3 interactions. Right inset shows key D1-D4 interactions.

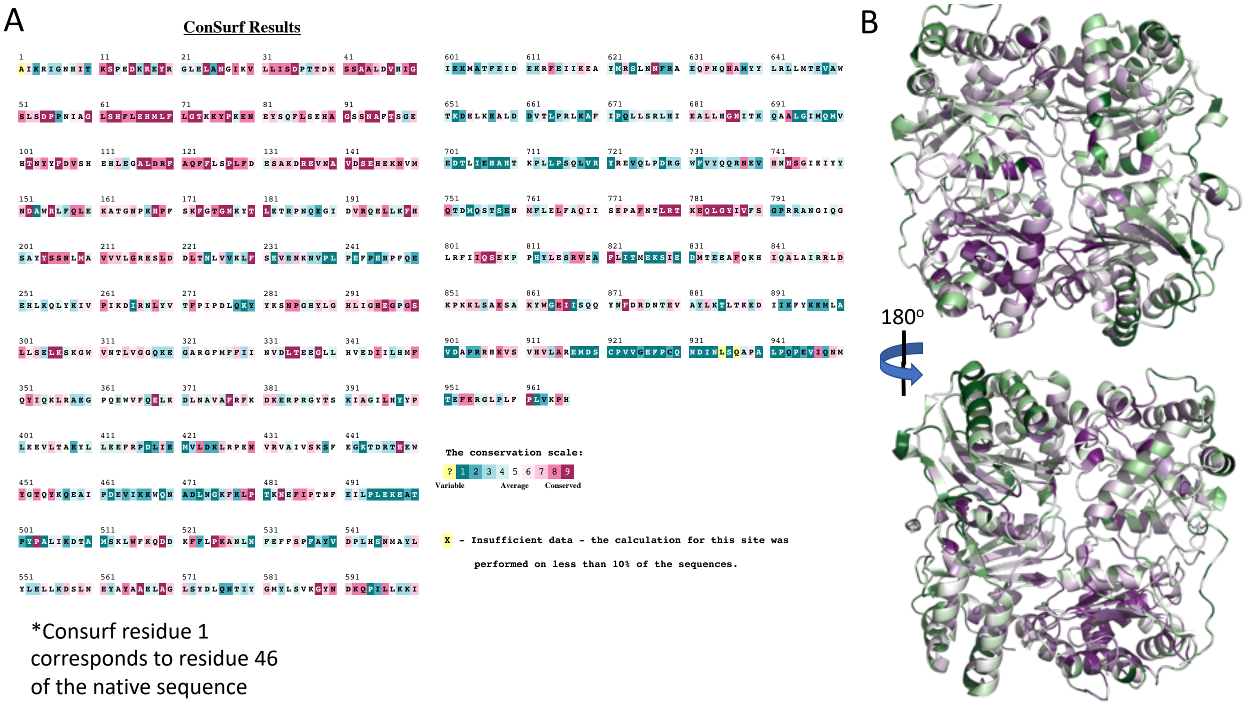


**Figure 6 – figure supplement 3 – IDE Consurf analysis. (A)** Conservation of IDE residues compared to known homologs. Residues colored on scale from conserved (purple) to variable (green). **(B)** Residue conservation results mapped onto the closed state IDE structure, residues colored as in **(A)**.

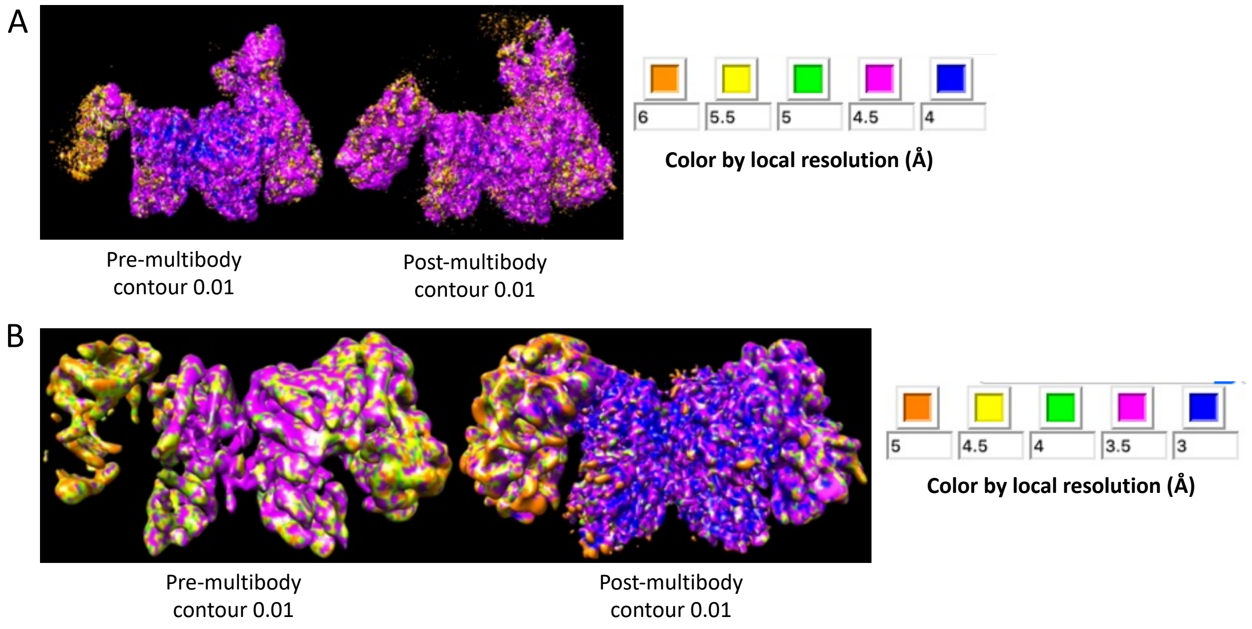


**Figure 2 – figure supplement 4 – Justification of multibody parameters. (A)** 3D volume of IDE with a single $Fab_v$ density before and after multibody refinement. **(B)** 3D volume of IDE with no $Fab_v$ density before and after multibody refinement. Volumes colored by local resolution, as indicated. The quality of the multibody results was assessed based on the subsequent improvement in the Coulomb potential map quality and calculated resolution. In the absence of any $F_v$ density, the maps resulting from multibody refinement had the best calculated resolution, yet the map quality was quite poor overall; much of the density appears globular and featureless, particularly in the IDE-N regions. Conversely, when $F_v$ density was present on both IDE-N bodies, the resolution of the multibody output maps (4.5 Å) was worse than the resolution of the input map (4.3 Å). Thus, we found that the greatest improvement occurred when the $F_v$ density was present on only the exterior body of the pO subunit. In this case, multibody refinement improved both the calculated resolution and density quality. See Methods for details on experimental setup.

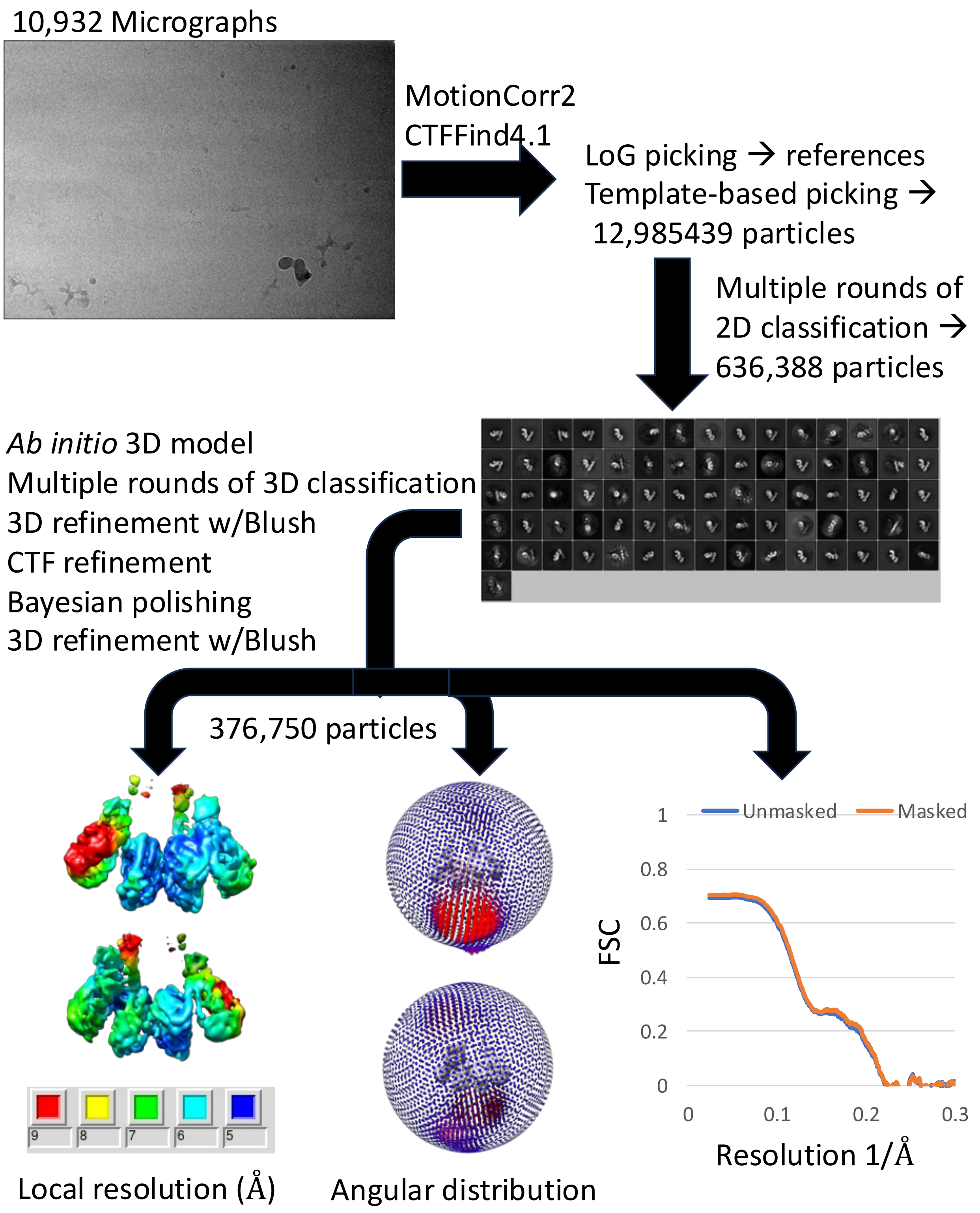

10,932 Micrographs
MotionCorr2
CTFFind4.1
LoG picking → references
Template-based picking →
12,985439 particles
Multiple rounds of
2D classification →
636,388 particles
*Ab initio* 3D model
Multiple rounds of 3D classification
3D refinement w/Blush
CTF refinement
Bayesian polishing
3D refinement w/Blush
376,750 particles
9
8
7
6
5
Local resolution (Å)
Angular distribution
Unmasked
Masked
1
0.8
0.6
0.4
0.2
0
FSC
0
0.1
0.2
0.3
Resolution 1/Å

**Figure 7 – figure supplement 1 - Processing info for time-resolved cryoEM analysis of IDE that was mixed with insulin only 123 milliseconds.**

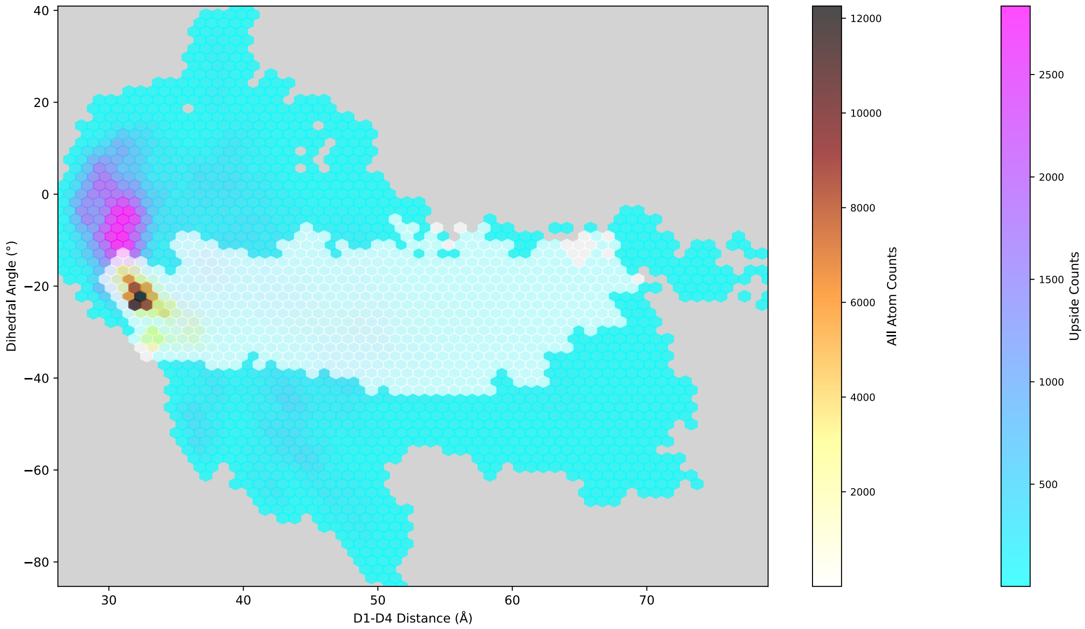


**Figure 8 – figure supplement 1 – Upside sampling of IDE conformational space.** The D1-D2-D3-D4 centers-of-mass dihedral angle and D1-D4 centers-of-mass distance were calculated for both subunits of IDE for every frame of our Upside simulations to represent the conformational space sampled during our simulations. Upside conformational space (heat map from cyan to magenta) is compared to the conformational space sampled in our all-atom MD simulations (heat map from white to brown, from figure 4 figure supplement 1). Each count corresponds to a single frame subunit measurement.

**Figure 3 - video 1 – Multibody translational component of motion.** Multibody analysis reveals major components of structural heterogeneity can be described by IDE-N swinging towards or away from IDE-C about a hinge formed by the interdomain linker region. Motion between the extreme start and end states of a representative eigenvector of structural heterogeneity. Cartoon model was fit into the start and end states and used to depict motion, monomer shown for clarity.

**Figure 3 – video 2 – Multibody rotational component of motion.** Multibody analysis reveals major components of structural heterogeneity can be described by IDE-N grinding against IDE-C like a screw. Motion between the extreme start and end states of a representative eigenvector of structural heterogeneity. Cartoon model was fit into the start and end states and used to depict motion, monomer shown for clarity.

**Figure 3 – video 3 – Normal mode analysis suggests a rotational component of motion.** The lowest frequency mode derived from normal mode analysis is consistent with the major rotational component of structural heterogeneity revealed by multibody analysis.

**Figure 3 – video 4 – Normal mode analysis suggests a translational component of motion.** The second lowest frequency mode derived from normal mode analysis is consistent with the major translational component of structural heterogeneity revealed by multibody analysis.

**Figure 8 – video 1 – Example Upside simulation of IDE in complex with insulin**. IDE and insulin colored as in Fig. 2. Plot and stills in Fig. 8 are derived from this trajectory.

| **Supplementary file 1: Cryo-EM data collection, refinement, and validation statistics of IDE for the 2:1 molar ratio of IDE to insulin.** | | | | | |
|---|---|---|---|---|---|
| **Data collection and processing** | | | | | |
| Microscope | Titan Krios | | | | |
| Camera | Gatan K3 | | | | |
| Automation software | Leginon | | | | |
| Magnification | 81,000 | | | | |
| Voltage (kV) | 300 | | | | |
| Frames collected per micrograph | 50 | | | | |
| Dose per frame (e-/Å2) | 1.36 | | | | |
| Total electron dose (e-/Å2) | 67.9 | | | | |
| Defocus range (µM) | 0.7 to 1.5 | | | | |
| Total micrographs | 7,611 | | | | |
| Initial particle images (no.) | 7,206,464 | | | | |
| | **O/O state** | **O/pO state** | **pO/pC state** | **O/pC state** | **pC/pC state** |
| Pixel size (Å) | 1.0842 | 1.0842 | 1.0842 | 1.0842 | 1.0842 |
| Final particle images (no.) | 77,973 | 328,870 | 76,379 | 304,011 | 1,341,061 |
| Symmetry imposed | C1 | C1 | C1 | C1 | C1 |
| Map resolution (Å) | 3.8 | 4.1 | 3.3 | 3.4 | 3.0 |
| FSC threshold | 0.143 | 0.143 | 0.143 | 0.143 | 0.143 |
| EMDB | EMD-24760 | EMD-24759 | EMD-24757 | EMD-24758 | EMD-24761 |
| **Refinement** | | | | | |
| Model resolution | | | | | |
| FSC 0.5 | 7.4 (7.7)[a] | 4.5 (7.4)[a] | 3.5 (3.8)[a] | 3.6 (3.9)[a] | 3.1 (3.2)[a] |
| FSC 0.143 | 3.6 (3.8)[a] | 3.4 (3.8)[a] | 3.1 (3.2)[a] | 3.3 (3.3)[a] | 3.0 (3.0)[a] |
| Sharpening B factor | -73.0 | -70.0 | -87.1 | -47.7 | -109.8 |
| Refinement package | PHENIX & COOT | PHENIX & COOT | PHENIX & COOT | PHENIX & COOT | PHENIX & COOT |
| Model composition | | | | | |
| Protein residues | 1867 | 1888 | 1898 | 1902 | 1926 |
| Total atoms | 15300 | 15465 | 15545 | 15584 | 15775 |
| B factors | | | | | |
| Protein | 64.38 | 129.58 | 70.19 | 80.87 | 26.93 |
| RMS deviations | | | | | |
| Bond length | 0.006 | 0.006 | 0.005 | 0.005 | 0.005 |
| Bond angle | 1.148 | 1.126 | 1.031 | 1.062 | 0.985 |
| Ramachandran (%) | | | | | |
| Favored | 96.99 | 94.36 | 96.34 | 95.92 | 97.07 |
| Allowed | 3.01 | 5.64 | 3.66 | 4.08 | 2.93 |
| Outliers | 0 | 0 | 0 | 0 | 0 |
| **Validation** | | | | | |

| | | | | | |
|---|---|---|---|---|---|
| MolProbity score | 1.51 | 1.67 | 1.34 | 1.49 | 1.3 |
| Poor rotamers (%) | 0.24 | 0.06 | 0.18 | 0.12 | 0.23 |
| Clash score | 6.07 | 5.26 | 3.03 | 4.25 | 3.44 |
| Correlation coefficient | 0.68 | 0.68 | 0.77 | 0.74 | 0.79 |
| Cbeta outliers | 0.11 | 0 | 0 | 0.06 | 0.05 |
| CaBLAM outliers | 1.89 | 2.73 | 1.76 | 1.86 | 1.9 |
| EMRinger score | 0.51 | 1.27 | 2.52 | 2.28 | 2.92 |
| PDB ID | 7RZH | 7RZG | 7RZE | 7RZF | 7RZI |

[a]Unmasked resolution is given in parentheses

| **Supplementary file 2: Time-resolved cryo-EM data collection, refinement, and validation statistics of IDE that was rapidly mixed with 5 fold molar excess of insulin for 123 milliseconds before vitrification** | |
|---|---|
| **Data collection and processing** | |
| Microscope | Titan Krios |
| Camera | Gatan K3 |
| Automation software | Leginon |
| Magnification | 81,000 |
| Voltage (kV) | 300 |
| Frames collected per micrograph | 50 |
| Dose per frame (e-/Å2) | 1.5 |
| Total electron dose (e-/Å2) | 65 |
| Defocus range (μM) | -0.7 to -2.5 |
| Total micrographs | 10,922 |
| Initial particle images (no.) | 12,985,439 |
| | **O/O state** |
| Pixel size (Å) | 1.06 |
| Energy filter | 20 eV slit |
| Final particle images (no.) | 376,750 |
| Symmetry imposed | C1 |
| Map resolution (Å) | 5.15 |
| FSC threshold | 0.143 |
| EMDB | EMD-72393 |
| **Refinement** | |
| Model resolution | |
| FSC 0.5 | 7.2 (7.2)[a] |
| FSC 0.143 | 4.9 (4.9)[a] |
| Sharpening B factor | -270.4 |
| Refinement package | PHENIX & COOT |
| Model composition | |
| Protein residues | 1888 |
| Total atoms | 15412 |
| B factors | |
| Protein | 258.04 |
| RMS deviations | |
| Bond length | 0.003 |
| Bond angle | 0.755 |
| Ramachandran (%) | |
| Favored | 94.36 |
| Allowed | 5.54 |
| Outliers | 0.11 |

| **Validation** | |
|---|---|
| MolProbity score | 2.18 |
| Poor rotamers (%) | 0.06 |
| Clash score | 19.85 |
| Cbeta outliers | 0 |
| CaBLAM outliers | 2.84 |
| EMRinger score | 0.26 |
| PDB ID | 9Y0H |
| [a]Unmasked resolution is given in parentheses | |

**Supplementary file 3 – Summary of multibody measurements for the Apo O/pO state**

| Component vector | Variance described (%) | Change in O state D1-D4 COM distance (Å) | Change in O state D1-D2-D3-D4 dihedral (degrees) | Change in pO state D1-D4 COM distance (Å) | Change in pO state D1-D2-D3-D4 dihedral (degrees) |
|---|---|---|---|---|---|
| 1 | 12.59 | -4.1 | 20.2 | -1.3 | -19.2 |
| 2 | 11.22 | 4.1 | -2.9 | -10.4 | 14.2 |
| 3 | 10.23 | -21.2 | -8.8 | 5.5 | 3.1 |
| 4 | 7.77 | 17.4 | -33.9 | -1.6 | -1.2 |
| 5 | 7.39 | 12.0 | 2.8 | 5.3 | 6.6 |
| 6 | 6.97 | -5.8 | -9.9 | 2.6 | -7.7 |
| 7 | 6.71 | -16.3 | -26.0 | -0.4 | 5.7 |
| 8 | 6.20 | -3.3 | -0.7 | 2.3 | -11.5 |
| 9 | 5.73 | 2.1 | -3.4 | -0.3 | 1.8 |

**Supplementary file 4 – Summary of multibody measurements for the IDE-ins (2:1) O/pO state**

| Component vector | Variance described (%) | Change in O state D1-D4 COM distance (Å) | Change in O state D1-D2-D3-D4 dihedral (degrees) | Change in pO state D1-D4 COM distance (Å) | Change in pO state D1-D2-D3-D4 dihedral (degrees) |
|---|---|---|---|---|---|
| 1 | 16.36 | -1.5 | 5.8 | 3.7 | -7.6 |
| 2 | 15.45 | -1.3 | -22.6 | 1.1 | -4.0 |
| 3 | 12.12 | 14.3 | -6.1 | 0.2 | 4.4 |
| 4 | 11.89 | -7.0 | 3.5 | 0.3 | 10.9 |
| 5 | 7.37 | -1.5 | -4.8 | -0.1 | 0.8 |
| 6 | 6.25 | 1.0 | 14.0 | -0.7 | -11.0 |
| 7 | 5.85 | 0.1 | 0.6 | 3.0 | -2.0 |
| 8 | 4.95 | -1.4 | 1.2 | 1.0 | -2.1 |
| 9 | 4.19 | 2.8 | 5.8 | -0.5 | -6.0 |

**Supplementary file 5 – Summary of multibody measurements for the IDE-ins (2:1) O/O state**

| Component vector | Variance described (%) | Change in $O_1$ state D1-D4 COM distance (Å) | Change in $O_1$ state D1-D2-D3-D4 dihedral (degrees) | Change in $O_2$ state D1-D4 COM distance (Å) | Change in $O_2$ state D1-D2-D3-D4 dihedral (degrees) |
|---|---|---|---|---|---|
| 1 | 17.57 | 5.3 | -7.4 | 0.4 | 1.4 |
| 2 | 15.88 | -18 | -27.8 | -0.8 | -5.3 |
| 3 | 13.00 | 0.0 | 4.6 | -12.2 | -16.5 |
| 4 | 11.73 | 0.1 | 2.4 | 15.0 | -21.1 |
| 5 | 8.52 | -0.7 | -11.2 | -2.5 | 11.4 |
| 6 | 8.21 | -0.3 | -10.8 | 5.7 | 9.0 |
| 7 | 7.12 | 5.2 | -0.7 | -1.9 | 1.0 |
| 8 | 5.45 | -18.0 | -16.3 | -1.2 | 13.1 |
| 9 | 3.20 | -2.5 | 3.0 | 7.3 | -4.3 |

**Supplementary file 6 – Summary of multibody measurements for the IDE-ins (2:1) O/pC state**

| Component vector | Variance described (%) | Change in O state D1-D4 COM distance (Å) | Change in O state D1-D2-D3-D4 dihedral (degrees) | Change in pC state D1-D4 COM distance (Å) | Change in pC state D1-D2-D3-D4 dihedral (degrees) |
|---|---|---|---|---|---|
| 1 | 15.77 | 6.8 | -15.6 | -3.4 | 1.2 |
| 2 | 13.79 | 0.1 | -4.8 | 5.4 | -20.3 |
| 3 | 12.05 | -0.1 | -1.9 | 11.4 | 0.9 |
| 4 | 11.19 | 1.8 | 22.6 | -0.2 | -5.1 |
| 5 | 10.09 | -0.6 | -8.8 | 1.7 | 12.2 |
| 6 | 7.91 | 0.1 | 1.9 | 0.6 | -10.9 |
| 7 | 7.04 | 2.3 | -9.9 | -0.8 | 0.8 |
| 8 | 6.29 | 5.6 | -2.5 | -2.2 | 3.4 |
| 9 | 5.07 | -0.8 | 3.9 | 4.9 | -6.3 |

**Supplementary file 7 – Summary of multibody measurements for the IDE-ins (2:1) pC/pC state**

| Component vector | Variance described (%) | Change in $pC_1$ state D1-D4 COM distance (Å) | Change in $pC_1$ state D1-D2-D3-D4 dihedral (degrees) | Change in $pC_2$ state D1-D4 COM distance (Å) | Change in $pC_2$ state D1-D2-D3-D4 dihedral (degrees) |
|---|---|---|---|---|---|
| 1 | 17.09 | 7.0 | -16.0 | -1.7 | -0.8 |
| 2 | 13.87 | 1.6 | 19.3 | 0.7 | 3.3 |
| 3 | 13.57 | 0.1 | 5.3 | -6.7 | -8.2 |
| 4 | 10.84 | -0.2 | 1.0 | 9.8 | -13.4 |
| 5 | 8.16 | -2.7 | -10.7 | 2.8 | 13.1 |
| 6 | 8.06 | 4.1 | -7.5 | 0.3 | 1.0 |
| 7 | 6.49 | 2.1 | -2.8 | -2.5 | 11.6 |
| 8 | 4.81 | 1.1 | -10.1 | -1.0 | 1.4 |
| 9 | 3.69 | -2.5 | 3.4 | 5.6 | -6.6 |

**Supplementary file 8 – Summary of multibody measurements for the IDE-ins (2:1) pO/pC state**

| Component vector | Variance described (%) | Change in pO state D1-D4 COM distance (Å) | Change in pO state D1-D2-D3-D4 dihedral (degrees) | Change in pC state D1-D4 COM distance (Å) | Change in pC state D1-D2-D3-D4 dihedral (degrees) |
|---|---|---|---|---|---|
| 1 | 24.94 | 4.6 | -9.1 | 0.6 | -0.3 |
| 2 | 16.96 | -0.1 | 1.9 | 8.2 | -11.4 |
| 3 | 12.5 | -1.0 | -10.9 | 1.6 | -1.5 |
| 4 | 8.92 | 0.7 | -0.2 | 2.1 | 6.2 |
| 5 | 8.54 | -1.2 | 3.5 | 1.7 | 3.0 |
| 6 | 6.15 | -0.1 | -1.4 | 0.0 | -5.0 |
| 7 | 3.98 | 0.4 | 7.4 | -1.0 | -7.6 |
| 8 | 2.79 | 1.1 | -2.9 | -1.0 | -0.6 |
| 9 | 2.31 | -2.3 | 2 | 3.1 | -3.7 |
